# Interneuronal dynamics facilitate the initiation of cortical spreading depression

**DOI:** 10.1101/2021.04.25.441350

**Authors:** Wolfgang Stein, Allison L. Harris

**Affiliations:** School of Biological Sciences, Illinois State University, Normal, IL 61790 USA; Department of Physics, Illinois State University, Normal, IL 61790 USA

**Keywords:** migraine, spike block, sodium channel inactivation, Familial Hemiplegic Migraines, pyramidal cell, GABAergic inhibition

## Abstract

Cortical spreading depression (CSD) is thought to precede migraine attacks with aura and is characterized by a slowly traveling wave of inactivity through cortical pyramidal cells. During CSD, pyramidal cells experience hyperexcitation with rapidly increasing firing rates, major changes in electrochemistry, and ultimately spike block that propagates slowly across the cortex. While the identifying characteristic of CSD is the pyramidal cell hyperexcitation and subsequent spike block, it is currently unknown how the dynamics of the cortical microcircuits and inhibitory interneurons affect the initiation of CSD.

We tested the contribution of cortical inhibitory interneurons to the initiation of spike block using a cortical microcircuit model that takes into account changes in ion concentrations that result from neuronal firing. Our results show that interneuronal inhibition provides a wider dynamic range to the circuit and generally improves stability against spike block.

Despite these beneficial effects, strong interneuronal firing contributed to rapidly changing extracellular ion concentrations, which facilitated hyperexcitation and led to spike block first in the interneuron and then in the pyramidal cell. In all cases, a loss of interneuronal firing triggered pyramidal cell spike block. However, preventing interneuronal spike block was insufficient to rescue the pyramidal cell from spike block. Our data thus demonstrate that while the role of interneurons in cortical microcircuits is complex, they are critical to the initiation of pyramidal cell spike block and CSD. We discuss the implications that localized effects on cortical interneurons have beyond the isolated microcircuit.

## 1 Introduction

Migraine is a disease afflicting more than 39 million men, women, and children in the U.S. and an estimated 1 billion people worldwide (Migraine Research Foundation, 2021). The migraine event is often accompanied by extreme sensitivity to light and sound that can last for hours to days. Some forms of migraines are preceded by auras, including visual or other disturbances, such as flashes of light, blind spots, tingling on one side of the face, arm, or leg, and difficulty speaking (Markus A. Dahlem, 2013; Dalkara & Moskowitz, 2017).

Migraine is thought to be preceded, and perhaps initiated, by cortical spreading depression (CSD), a pathology characterized by a slowly traveling wave of inactivity in cortical pyramidal cells (C. Ayata, 2009; Miura et al., 2007; Wei et al., 2014; Zandt et al., 2015). It is not currently known what initiates CSD, but the characteristic indicators are a rapid increase in firing frequency of the pyramidal cells followed by a loss of spiking that then propagates slowly through the cortex (C. Ayata, 2009; Herreras et al., 1994). The propensity for pyramidal spike block increases with increased excitability of the cortical circuits, and the loss of spiking in pyramidal cells results from a continued depolarization that leads to sodium channel inactivation and spike block (Wei et al., 2014; Zandt et al., 2015).

Because the spreading inactivity wave is a collective behavior of a large-scale network, its underlying mechanisms remain elusive. One proposed cause of the increased excitation in the pyramidal cells is the loss of potassium homeostasis in the extracellular medium during heightened activity of the pyramidal cells and the local inhibitory interneurons. Several lines of evidence from both biological experiments and computational models have provided support for this mechanism and demonstrated that spike block in pyramidal cells is associated with increased extracellular potassium concentration that can be initiated experimentally (Cenk Ayata & Lauritzen, 2015; Chronicle et al., 2006; M. A. Dahlem & Chronicle, 2004).

However, CSD may also be facilitated by many factors that increase neuronal excitability, including swelling of the cells or mutations in ion channels and pumps (Cenk Ayata & Lauritzen, 2015; Markus A. Dahlem et al., 2014; Dalkara & Moskowitz, 2017; Hübel et al., 2017; Tuttle et al., 2019; Ullah et al., 2015; Wei et al., 2014).

In particular, a set of known genetic mutations has been identified in a small subset of migraineurs. Collectively, these genetic mutations lead to disorders known as Familial Hemiplegic Migraines (Dalkara & Moskowitz, 2017; De Fusco et al., 2003; Dichgans et al., 2005; Ophoff et al., 1996), which can be caused by gain-of-function mutations for voltage gated ion channels or loss-of-function mutations in the sodium-potassium (Na/K) pump. More recently, mutations in the sodium channel prevalent in cortical interneurons have been implicated in the initiation of CSD. Specifically, a mutation of the Na_V_ 1.1 sodium channel (SCN1A gene) has been identified as FHM-3 (Dichgans et al., 2005; Mantegazza & Broccoli, 2019; Tiwari et al., 2020). At this point, it is not clear whether this mutation causes a gain or loss of function in the channel (Cestèle et al., 2008, 2013; Kahlig et al., 2008; Mantegazza & Broccoli, 2019). In epileptic pathologies, a mutation in the Na_V_ 1.1 channel causes a loss of function in the channel leading to reduced excitation of cortical interneurons, and a subsequent disinhibition and hyperexcitability of the pyramidal cells (Hedrich et al., 2014). Conversely, evidence in cell cultures points to a gain of function in the channel, leading to hyperexcitability of the interneurons (Mantegazza & Broccoli, 2019). Despite these conflicting hypotheses regarding the role of the FHM-3 mutation in spike block initiation, it is clear that the interneurons are critical in this process, a notion also supported by computational studies (Desroches et al., 2019). However, the specific mechanisms and contributions of the cortical interneurons in the CSD process are still not well understood.

To better understand the cellular properties and mechanisms leading to CSD, simplified cortical microcircuits can be used to study the initiation of spike block in pyramidal cells, which serves as an indicator of CSD. Previous studies have suggested that strong excitatory input to pyramidal cells or interneurons can lead to increased potassium leak that ultimately drives the pyramidal cells into a sodium block (Dalkara & Moskowitz, 2017; Wei et al., 2014). This finding is not surprising given that potassium accumulates quickly in tight extracellular spaces, and that homeostatic responses of the Na/K pump are comparatively slow. Previous models also indicated that in conditions with heightened interneuron excitability, the dynamic range of pyramidal cell firing is small, with weakly firing pyramidal cells already eliciting sodium block (Desroches et al., 2019). This suggests that increased interneuronal excitability decreases stability in the negative feedback loop within the cortical microcircuit. However, the influence of extracellular potassium accumulation on the interneurons themselves has not been studied, which limits our understanding of the full dynamic range of the cortical microcircuits. Additionally, the trigger for the initial surge in neuron firing that leads to the sodium block remains unidentified. While it is clear that extracellular potassium mediates the initiation, it is unknown if the increased firing of the pyramidal cell or that of the interneuron destabilizes the system. It is also unclear what role circuit properties play in the destabilization of the cortical network.

The major hypothesis driving this study is that heightened interneuronal excitability stabilizes cortical microcircuits and diminishes spike block in interneurons. We further hypothesize that if interneuronal inhibition is lost, either due to a loss of excitation or interneuronal spike block, it will lead to spike block in pyramidal cells and depression of the entire circuit. We test our hypothesis in a simplified cortical circuit model consisting of a negative feedback loop between a pyramidal cell and an interneuron. Ion concentrations in the extra- and intracellular space are included in the model and we differentiate the effects of interneuronal firing frequency from extracellular ion concentrations on the initiation of spike block.

## 2 Methods

We use two separate but similar models to examine the effects of ion concentrations on the interneuron and the reciprocal effects of the interneuron on extracellular ion concentrations. Both models are 2-cell computational models consisting of an interneuron and pyramidal cell. The neurons interact through a feedback loop, with reciprocal inhibition between the two neurons, as shown in Fig 1A. The 2-cell, Pyramidal cell, Interneuron (2PI) model includes the effects of ion concentrations on the interneuron, as well as the interneuron’s effect on the extracellular ion concentrations. The 2-cell, Pyramidal cell, Interneuron minus ion Concentration (2PI-C) model is similar, but has fixed interneuron intracellular ion concentrations and the interneuron’s effect on the extracellular ion concentrations is only included through its voltage gated potassium current. Both models are described in detail below.

**Fig. 1.**
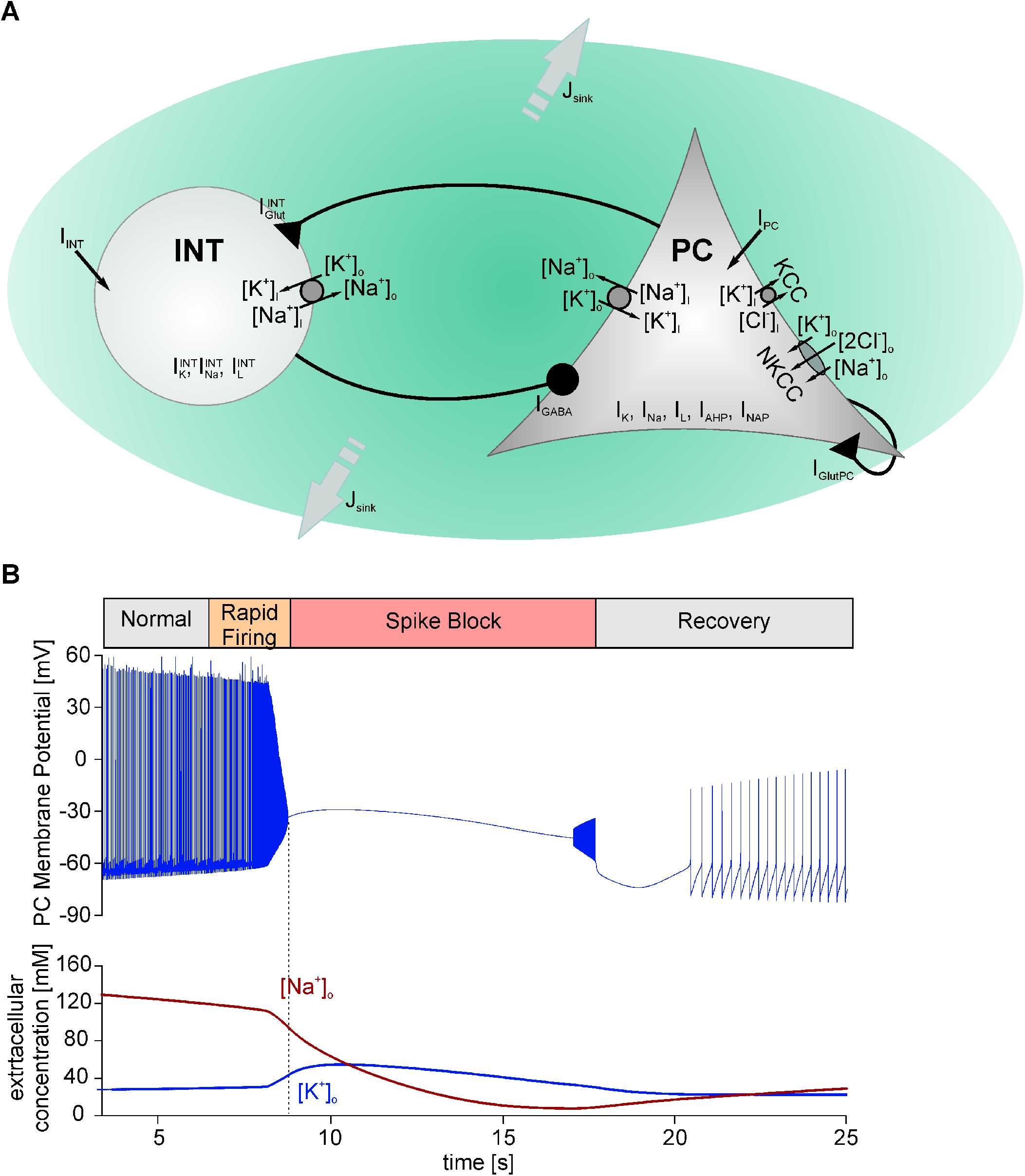
A. Schematic of the cortical two-cell microcircuit model. The pyramidal cell and the interneuron contained a sodium-potassium pump 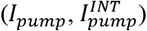, as well as leak 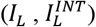, fast sodium 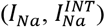 and delayed rectifier potassium currents 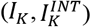. The pyramidal cell additionally contained a calcium-dependent potassium current (*I_AHP_*), a persistent sodium current (*I_Nap_*), a potassium/chloride cotransporter (*J*_*KCC*2_) and a sodium-potassium-chloride cotransporter (*J_NKCC_*). The pyramidal cell excited the interneuron via a glutamatergic synapse 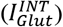, and the interneuron inhibited the pyramidal cell via a GABA-ergic synapse (*I_GABA_*). In addition, the pyramidal cell possessed an autoexcitatory glutamatergic synapse to account for the excitatory influence of other pyramidal cells (*I_GlutPC_*). Ion concentration changes caused by ion flow through channels and pumps allowed for ion accumulation in the extracellular space, which was also equipped with a potassium sink to account for diffusion and removal of extracellular potassium. Both neurons could be injected with depolarizing currents (*I_INT_* and *I_PC_*, respectively) to change their excitability state. B. Example spike block in a pyramidal cell. Network parameters were: *I_pc_* = 5 μA/cm^2^., *I_INT_* = 1 μA/cm^2^, *g_GABA_* = 0.4 mS/cm^2^. Top: Spike block is characterized by a phase of rapidly increasing firing frequencies, followed by the complete loss of action potentials. Bottom: Changes in extracellular sodium (red) and potassium (blue) concentrations that accompany spike block.

### 2.1 2PI Model

#### 2.1.1 Pyramidal Cell

The pyramidal cell is modeled as a single compartment model containing voltage gated sodium channels *I_Na_*, voltage gated potassium channels *I_K_*, calcium activated potassium channels *I_AHP_*, leak channels *I_L_*, and persistent sodium channels *I_Nap_*. In addition, the pyramidal cell model contains a sodium-potassium pump *I_pump_*, an injected current *I_pc_*, and two synaptic currents *I_GABA_* and *I_glut_pc__*. The inhibitory synapse from the interneuron is *I_GABA_*, and *I_glut_pc__* is a self-excitation used to represent the influence of additional cortical pyramidal cells. Then, the membrane potential of the pyramidal cell *V_pc_* is given by the modified Hodgkin-Huxley equations (Hodgkin & Huxley, 1952)

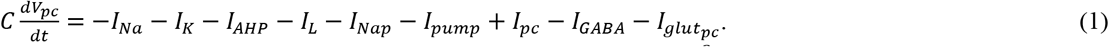

The individual voltage-gated current densities (in units of *μA*/*cm*^2^) are given by

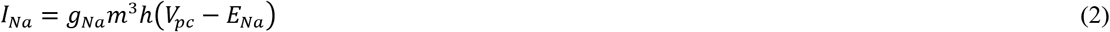

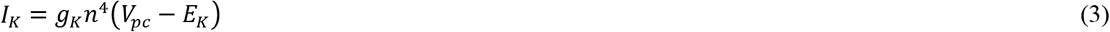

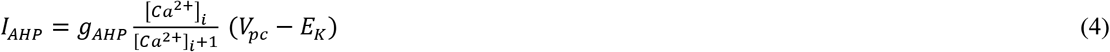

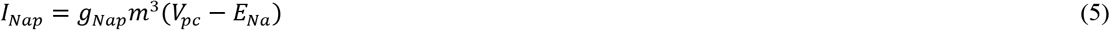

where *g_x_* (*x* = *Na, K, AHP, Nap*) is the maximum conductance and *m, h*, and *n* are the activation and inactivation variables representing the fraction of open and closed ion channel gates. All parameter values used for the pyramidal cell are listed in

Table 1. The equilibrium potentials *E_x_* (*x* = *Na, K, Cl*) vary with intra- and extracellular ion concentrations according to the Nernst equation

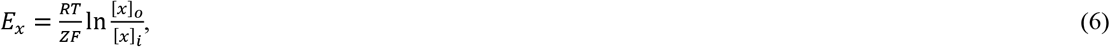

where *Z* is the charge of the ion, *R* is the universal gas constant, *T* is the temperature of the system, and *F* is the Faraday constant. The concentrations are denoted as [*x*]_*i,o*_, where *x* is the ion and *i, o* refers to intra-or extracellular respectively.

**Table 1.** Pyramidal cell parameters.

| Parameter | Value | Definition |
| --- | --- | --- |
| $C$ | $1 \mu F/cm^2$ | Pyramidal cell membrane capacitance (Wei et al., 2014) |
| $g_{Na}$ | $100 \text{ mS/cm}^2$ | Pyramidal cell maximum sodium conductance (Börger et al., 2005; Wei et al., 2014) |
| $g_K$ | $80 \text{ mS/cm}^2$ | Pyramidal cell maximum potassium conductance (Börger et al., 2005) |
| $g_{AHP}$ | $1.5 \text{ mS/cm}^2$ | Pyramidal cell maximum calcium dependent potassium conductance (Desroches et al., 2019) |
| $g_{Nap}$ | $1 \text{ mS/cm}^2$ | Pyramidal cell maximum persistent sodium conductance (Desroches et al., 2019) |
| $\rho_{pump}$ | $0.25 \text{ mM/s}$ | Sodium-Potassium pump strength (Desroches et al., 2019) |
| $\gamma$ | $0.044 \text{ mmol/(C cm)}$ | Conversion factor between $K^+$, $Na^+$, and $Cl^-$ current densities and rates of ion concentration change (Cressman et al., 2009) |
| $K_{sat}$ | $3.5 \text{ mM}$ | Extracellular potassium concentration when pump is at half capacity (Wei et al., 2014) |
| $Na_{sat}$ | $22 \text{ mM}$ | Pyramidal cell intracellular sodium concentration when pump is at half capacity (Desroches et al., 2019) |
| $I_{pc}$ | $\mu A/cm^2$, Variable | Pyramidal cell injected current density |
| $g_{GABA}$ | $\text{mS/cm}^2$, Variable | Maximum GABA-A conductance |
| $g_{glut}$ | $0.1 \text{ mS/cm}^2$ | Maximum glutamate conductance (Desroches et al., 2019) |
| $E_{glut}$ | $0 \text{ mV}$ | Equilibrium potential for glutamate synapse (Desroches et al., 2019) |
| $\tau_{GABA}$ | $9 \text{ ms}$ | Decay constant for the GABA synapse (Desroches et al., 2019) |
| $\tau_{glut_{pc}}$ | $3 \text{ ms}$ | Decay constant for the pyramidal cell self-excitatory synapse (Desroches et al., 2019) |
| $R$ | $8.3145 \text{ J/(mol K)}$ | Universal gas constant |
| $T$ | $310 \text{ K}$ | Temperature of the system (body temperature was used) |
| $F$ | $96485.3 \text{ A s/mol}$ | Faraday constant |
| $g_{K,L}$ | $0.05 \text{ mS/cm}^2$ | Pyramidal cell maximum potassium leak conductance (Wei et al., 2014) |
| $g_{Na,L}$ | $0.0015 \text{ mS/cm}^2$ | Pyramidal cell maximum sodium leak conductance (Desroches et al., 2019) |
| $g_{Cl,L}$ | $0.015 \text{ mS/cm}^2$ | Pyramidal cell maximum chloride leak conductance (Desroches et al., 2019) |
| $\phi$ | $1$ | Temperature factor for Hodgkin-Huxley gating variables (Börger et al., 2005) |

The leak current *I_L_* is composed of three terms, *I_K,L_, I_cl,L_*, and *I_Na,L_* representing the potassium, chloride, and sodium leak currents, respectively

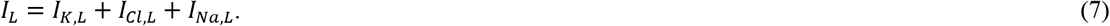

Each of the leak currents is given by an equation of the form (*x* = *K, Cl, Na*)

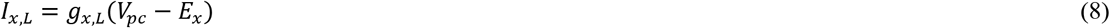

with maximum conductances *g_x,L_*.

The sodium-potassium pump serves to maintain the sodium and potassium concentration gradients by exporting sodium and importing potassium ions through the membrane. The current through the pump is modeled as a product of two sigmoidal functions dependent on each of the ion concentrations (Cressman et al., 2009)

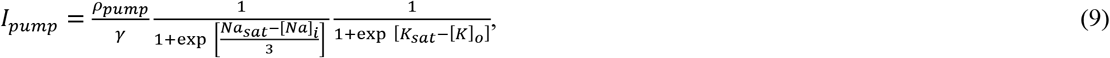

where *ρ_pump_* is the pump strength, *γ* is a conversion factor to convert from current density to rate of ion concentration change, and *K_sat_* and *Na_sat_* are the extracellular potassium and intracellular sodium concentrations at which the pump is operating at half its maximum activity.

The synaptic currents are given by (Desroches et al., 2019)

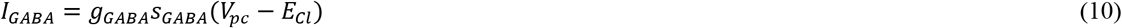

and

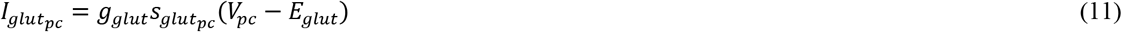

where *g_GABA_* and *g_glut_* are the maximum GABA-A and glutamate conductances, *E_cl_* is dependent upon the chloride concentration (see Eq. 6), and *E_glut_* is the equilibrium potential for the glutamate synapse. The quantities *s_GABA_* and *s_glut_pc__* are activation variables for the inhibitory synapse and the self-excitatory synapse that are set to unity following each spike of the interneuron or pyramidal cell, respectively (*V_INT,PC_* > 0). Following each spike, they are subsequently governed by equations of the form (*x* = *GABA, glut_pc_*)

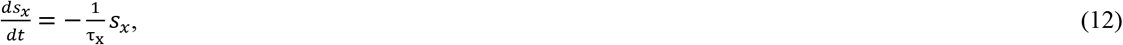

where *τ_x_* is the decay constant.

The activation and inactivation rate equations are (Börgers et al., 2005; Desroches et al., 2019)

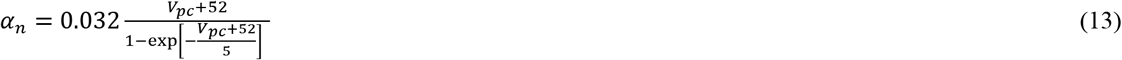

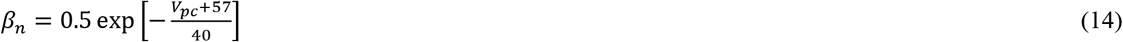

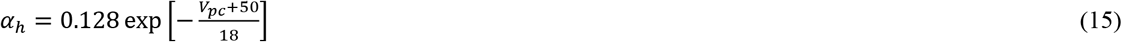

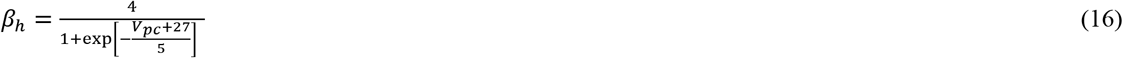

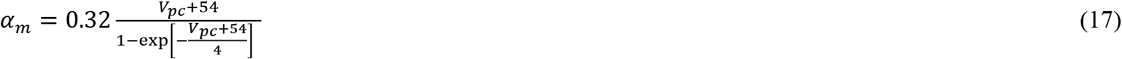

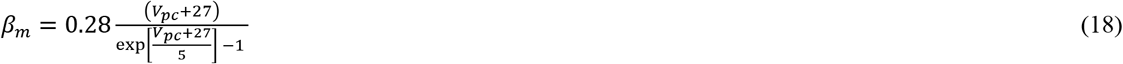

and the differential equations governing the activation gates are

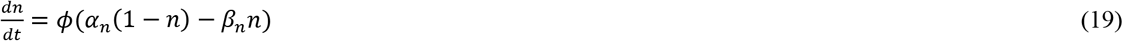

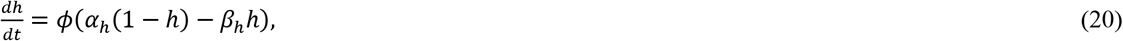

where *ϕ* is a temperature factor.

For the fast sodium channel, the steady state form of the sodium activation gate is used because the activation is assumed to be much faster than the change in voltage.

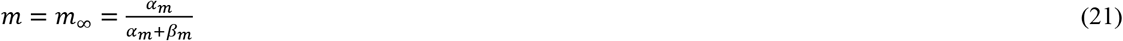

#### 2.1.2 Interneuron

Similar to the pyramidal cell, the interneuron was modeled as a single compartment model containing voltage gated sodium channels 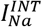, voltage gated potassium channels 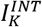, and leak channels 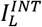. In addition, the interneuron model contained an injected current *I_INT_*, an excitatory synaptic current *I_glut_INT__*, and a sodium-potassium pump 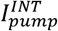. Then, the membrane potential of the interneuron *V_INT_* is given by the modified Hodgkin-Huxley equations (Hodgkin & Huxley, 1952)

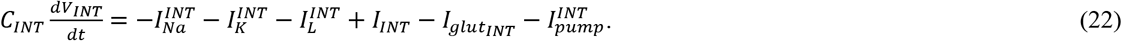

The individual voltage-gated current densities are given by

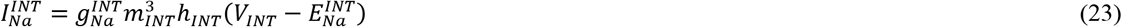

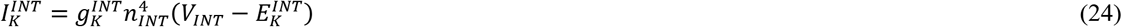

where 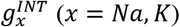 is the maximum conductance and *m_INT_, h_INT_*, and *n_INT_* are the activation and inactivation variables representing the fraction of open and closed ion channel gates. Parameter values used for the interneuron are listed in Table 2.

**Table 2.** Interneuron parameters.

| Parameter | Value | Definition |
| --- | --- | --- |
| $C_{INT}$ | $1 \mu F/cm^2$ | Interneuron membrane capacitance (Wang & Buzsáki, 1996) |
| $g_{Na}^{INT}$ | $35 \text{ mS/cm}^2$ | Interneuron maximum sodium conductance (Wang & Buzsáki, 1996) |
| $g_K^{INT}$ | $9 \text{ mS/cm}^2$ | Interneuron maximum potassium conductance (Wang & Buzsáki, 1996) |
| $g_{K,L}^{INT}$ | $0.08276 \text{ mS/cm}^2$ | Interneuron maximum potassium leak conductance<sup>1</sup> (Wang & Buzsáki, 1996) |
| $g_{Na,L}^{INT}$ | $0.0172 \text{ mS/cm}^2$ | Interneuron maximum sodium leak conductance<sup>1</sup> (Wang & Buzsáki, 1996) |
| $I_{INT}$ | $\mu A/cm^2$, Variable | Interneuron injected current density |
| $\gamma_I$ | $0.0286 \text{ mmol/(C cm)}$ | Conversion factor between current density and rate of ion concentration change (Desroches et al., 2019) |
| $\tau_{glut_{INT}}$ | $3 \text{ ms}$ | Decay constant for excitatory synapse on the interneuron (Desroches et al., 2019) |
| $\phi$ | $5$ | Temperature factor for Hodgkin-Huxley gating variables (Wang & Buzsáki, 1996) |

The equilibrium potentials 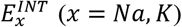 vary with intra- and extracellular ion concentrations according to Eq. (6) 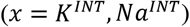.

The leak current 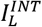 is composed of two terms, 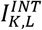 and 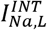 representing the potassium and sodium leak currents, respectively

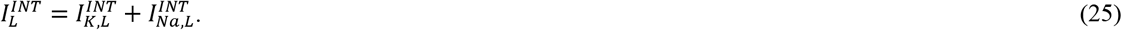

Each of the leak currents is given by an equation of the form (*x* = *K, Na*)

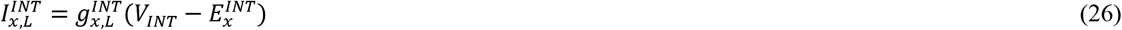

with maximum conductances 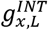.

The excitatory synaptic current is given by

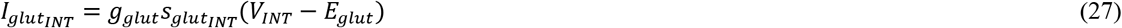

where *g_glut_* and *E_glut_* are the same as for the pyramidal cell. The quantity *s_glut_INT__* is the activation variable for the excitatory synapse and is set to unity following each spike of the pyramidal cell (*V_PC_* > 0). It is subsequently governed by Eq. (12) with *x* = *glut_INT_*.

The sodium-potassium pump current density is given by

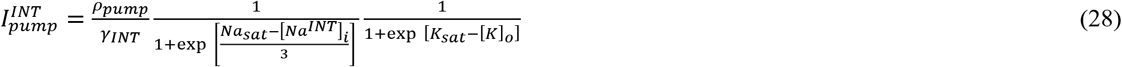

where *γ_INT_* is a conversion factor to convert from current density to rate of ion concentration change. The activation and inactivation rate equations are (Wang & Buzsáki, 1996)

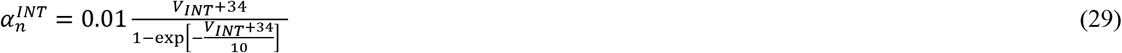

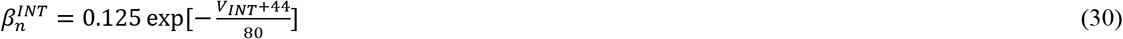

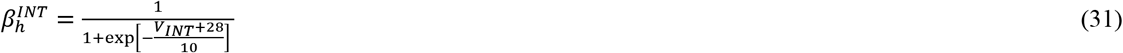

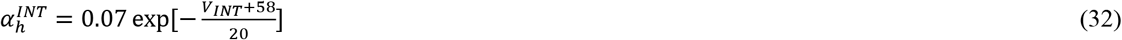

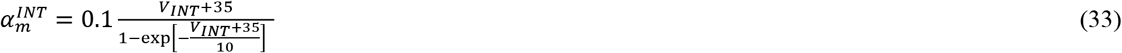

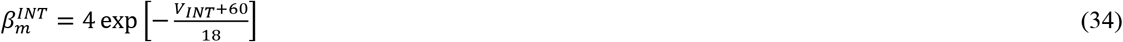

and the differential equations governing the activation gates are of the same form as Eqs. (19) and (20). We again use the steady state form of the activation of the sodium channel gates because the activation is assumed to be much faster than the change in voltage

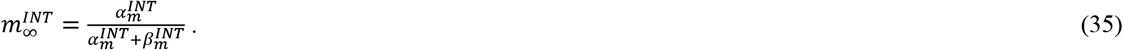

#### 2.1.3 Ion Concentrations

The intra- and extracellular ion concentrations are updated continuously in real time in the simulation and the relevant parameter values are given in Table 3.

**Table 3.** Ion concentration parameters.

| Variable | Value | Definition |
| --- | --- | --- |
| $\beta$ | 4 | Ratio of intracellular to extracellular volume (Syková & Nicholson, 2008) |
| $\tau$ | 1000 | Conversion from seconds to milliseconds (Desroches et al., 2019; Wei et al., 2014) |
| $\rho_{Kcc2}$ | 0.3 mM/s | Pyramidal cell maximum KCC2 cotransporter strength (Wei et al., 2014) |
| $\rho_{NKcc}$ | 0.1 mM/s | Pyramidal cell maximum NKCC1 cotransporter strength (Wei et al., 2014) |
| $\epsilon_K$ | 0.4 s<sup>-1</sup> | Rate factor for potassium loss to the surrounding environment (Desroches et al., 2019) |
| $K_{bath}$ | 3.5 mM | Potassium concentration of the surrounding environment (Desroches et al., 2019) |
| $g_{Ca}$ | 1 mS/cm<sup>2</sup> | Maximum calcium conductance (Wang, 1998) |
| $\epsilon_i$ | 0.002 mmol/(C cm) | Conversion factor between calcium current density and rate of ion concentration change (Wang, 1998) |
| $\tau_{ca}$ | 80 ms | Time constant for calcium buffering and extrusion (Helmchen et al., 1996; Markram et al., 1995; Svoboda et al., 1997; Wang, 1998) |

##### Potassium

In the extracellular space, the rate of change of potassium concentration is determined by the rate of potassium absorbed by the pyramidal cell 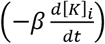 and the interneuron 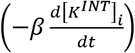, as well as a loss of potassium to the surrounding environment (−*J_sink_*)

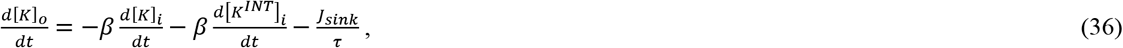

where *τ* is a conversion factor from seconds to milliseconds and *β* is the ratio of intracellular to extracellular volume.

The rate of potassium concentration change in the pyramidal cell is dependent upon the current densities from the potassium-dependent channels (*I_K_, I_AHP_, I_K,L_, I_pump_*), as well as concentration changes due to the K^+^/Cl^-^ cotransporter KCC2 (*J*_*Kcc*2_) (Payne et al., 2003) and the Na^+^/K^+^/2Cl^-^ cotransporter (*J_NKcc_*)

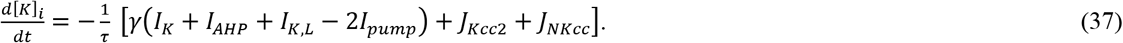

The cotransporter rates of concentration change are given by (Øyehaug et al., 2012)

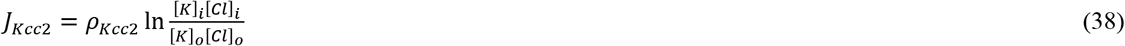

and

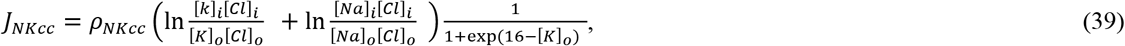

where *ρ*_*Kcc*2_ and *ρ_NKcc_* are the maximum KCC2 and NKCC1 cotransporter strengths.

The current density of the calcium-activated potassium channels *I_AHP_* depends on the pyramidal cell intracellular calcium concentration (Eq. (4)), which is modeled as a leaky integrator (Helmchen et al., 1996; Tank et al., 1995; Traub, 1982; Wang, 1998)

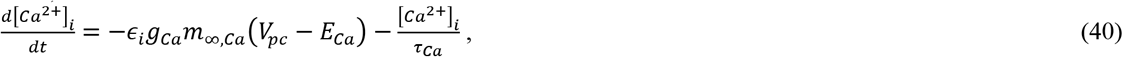

where *ϵ_i_* is a conversion factor between current density and rate of ion concentration change, *g_Ca_* is the maximal conductance, *τ_Ca_* is the time constant for calcium buffering and extrusion mechanisms, and *m_∞,Ca_* is the steady state representation of the calcium activation gates

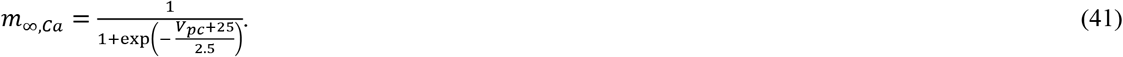

The rate of chloride concentration change in the extracellular space is found from the rate of change of intracellular chloride concentration in the pyramidal cell

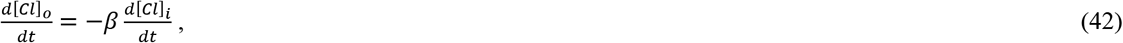

which is determined by the chloride-dependent current densities (*I_GABA_, I_Cl,L_*) and the concentration changes due to the cotransporters (*J*_*Kcc*2_, *J_NKcc_*)

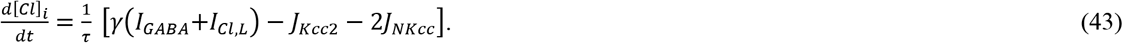

The factor of 2 multiplying *J_NKcc_* is indicative of the NKCC1 cotransporter transporting two chloride ions.

The rate of potassium concentration change in the interneuron is dependent upon the current densities from the potassium-dependent channels 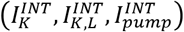

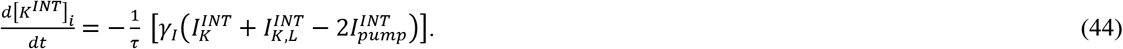

The factors of 2 multiplying *I_pump_* in Eq. (37) and 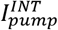 in Eq. (44) are indicative of the sodium-potassium pump transferring two potassium ions into the cell.

To model the loss of extracellular potassium to the surrounding environment, including through glial cells, an ion sink is included of the form

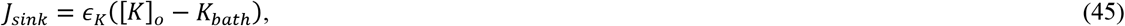

where *ϵ_K_* is the rate factor at which potassium is lost to the surrounding environment and *K_bath_* is the potassium concentration of the surrounding environment.

##### Sodium

The rate of change of sodium concentration in the extracellular space is also found by conservation of ions and is dependent upon the rate of sodium expelled from the pyramidal cell and the interneuron

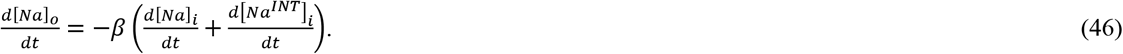

The rate of sodium concentration change in the pyramidal cell is dependent upon the current densities from the sodium-dependent channels (*I_Na_, I_Nap_, I_Na,L_, I_pump_*), as well as concentration changes due to the NKCC1 cotransporter (*J_NKcc_*)

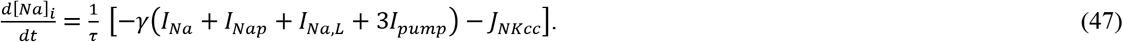

The rate of sodium concentration change in the interneuron was dependent upon the current densities from the sodium-dependent channels 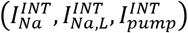

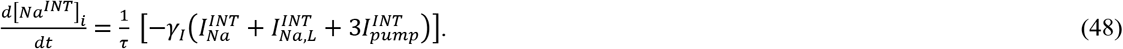

The factors of 3 multiplying *I_pump_* in Eq. (47) and 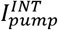 in Eq. (48) are indicative of the sodium-potassium pump expelling three sodium ions from the cell.

### 2.2 2PI-C Model

In the 2PI-C model, the K^+^/Na^+^ pump in the interneuron 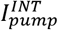 was excluded and the potassium and sodium equilibrium potentials did not depend on the ion concentrations, but instead were set to constant values 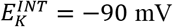 and 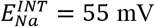. The interneuron intracellular potassium and sodium concentrations were not calculated and therefore Eqs. (44) and (48) don’t exist in the 2PI-C model. Also, conservation of ions in the 2PI-C model yields modified versions of Eqs. (36) and (46) for the extracellular potassium and sodium concentration

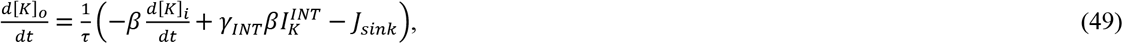

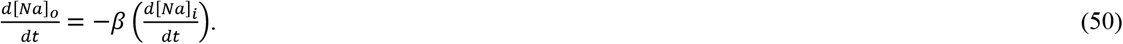

All other features of the 2PI-C model remained identical to the 2PI model.

### 2.3 Numerical Method

The simulations were run using a custom FORTRAN90 program that was parallelized to calculate multiple independent models using MPI. The differential equations were solved with a 4^th^ Order Runge-Kutta algorithm (William H. Press et al., 1992) with a timestep of 0.029 ms. The initial conditions for the integrated quantities are listed in Table 4. The 2PI model is available for download at ModelDB. Model source code is available at https://modeldb.yale.edu/267033 Use the password: steinharris

**Table 4.** Initial conditions for numerical calculations.

| Variable | Initial Condition | Description |
| --- | --- | --- |
| $V_{PC}$ | -70 mV | Pyramidal Cell membrane potential |
| $V_{INT}$ | -70 mV | Interneuron membrane potential |
| $n$ | 0 | Activation variable of the potassium current |
| $h$ | 1 | Inactivation variable of the sodium current |
| $m$ | 0 | Activation variable of the sodium current |
| $[Ca^{2+}]_i$ | 0 | Pyramidal cell intracellular calcium concentration |
| $[K]_i$ | 140 | Pyramidal cell intracellular potassium concentration |
| $[Na]_i$ | 12 | Pyramidal cell intracellular sodium concentration |
| $[Cl]_i$ | 5 | Pyramidal cell intracellular chloride concentration |
| $[K]_o$ | 5 | Extracellular potassium concentration |
| $[Na]_o$ | 140 | Extracellular sodium concentration |
| $[Cl]_o$ | 119 | Extracellular chloride concentration |
| $s_{GABA}$ | 0 | GABA-A synaptic variable |
| $s_{glut_{pc}}$ | 0 | Activation variable of the pyramidal cell self-excitatory glutamatergic synapse |
| $s_{glut_{INT}}$ | 0 | Activation variable of the pyramidal to interneuron glutamatergic synapse |
| $n_{INT}$ | 0 | Activation variable of the potassium current |
| $h_{INT}$ | 1 | Inactivation variable of the sodium current |
| $m_{INT}$ | 0 | Activation variable of the sodium current |
| $[K^{INT}]_i$ | 145.3 | Interneuron intracellular potassium concentration<sup>2</sup> |
| $[Na^{INT}]_i$ | 17.9 | Interneuron intracellular sodium concentration<sup>2</sup> |

## 3 Results

Fluctuating ion concentrations in cortical extracellular space have been indicated to elicit spike block in cortical pyramidal cells, particularly in states of high cortical excitability (Cenk Ayata & Lauritzen, 2015; Chronicle et al., 2006; M. A. Dahlem & Chronicle, 2004). However, pyramidal cells do not operate in isolation, but are embedded in a network of other pyramidal cells and interneurons. Ion concentrations change with spike activity in all of these neurons, making it important to assess the impact of all network neurons on ion concentrations, as well as the influence of ion concentrations on network neurons.

To address these issues, we modeled a cortical microcircuit consisting of one pyramidal cell and one interneuron, with the pyramidal cell exciting the interneuron via a glutamatergic synapse and the interneuron inhibiting the pyramidal cell via a GABA-ergic synapse (Figure 1A). In addition, to account for the presence of glutamatergic excitation from other pyramidal cells, a self-excitatory glutamate synapse was added to the pyramidal cell. Neuronal, synaptic, and model parameters for calculating ion concentrations were selected based on previously published models of cortical neurons (Tables 1 - 4, for details see Materials and Methods). While this microcircuit cannot represent the collective behavior of a larger cortical network, it can inform us of the cellular mechanisms that trigger spike block. The initiation of spike block serves as an indicator of instability in the network and is influenced by the excitation levels of both neurons, the inhibitory feedback synapse, and the extracellular ion concentrations. Figure 1B shows an example of spike block in a pyramidal cell that was induced by a high level of excitation of the pyramidal cell. The characteristic features that identify spike block are a ramping up of the firing frequency to extreme values of several hundred Hz, followed by a loss of spiking for an extended period of time during which the membrane potential remains depolarized. During this refractory period, the membrane potential initially continues to depolarize until it reaches −25 to −30 mV, after which a slow hyperpolarization starts. The pyramidal cell’s behavior coincided with changes in the concentrations of extracellular potassium and sodium. During the time that firing frequency ramped up, there was a slow accumulation of potassium, and a mild drop in sodium concentration. At the highest firing frequencies, immediately preceding spike block, both extracellular concentrations changed drastically, with potassium rising and sodium declining. Both concentrations then continued in an altered state for almost the full duration of the refractory period.

While the influence of changing ion concentrations on the pyramidal cells is well established (Cenk Ayata & Lauritzen, 2015; Chronicle et al., 2006; M. A. Dahlem & Chronicle, 2004), the effects on interneuronal activity and the resulting consequences for pyramidal cell activity are unclear. To examine these effects on the initiation of spike block in the cortical circuit, we compared two models. In one model (2PI-C), the ionic concentrations only affected the pyramidal cell, while in the other model (2PI), both neurons were affected.

In both models, the pyramidal cell contained a sodium-potassium pump (*I_pump_*), leak (*I_L_*), fast sodium (*I_Na_*), delayed rectifier potassium currents (*I_K_*), calcium-dependent potassium current (*I_AHP_*), persistent sodium current (*I_Nap_*), potassium/chloride cotransporter (*J_KCC2_*) and sodium-potassium-chloride cotransporter (*J_NKCC_*). The interneuron was simulated with a comparatively simpler model, with only leak 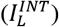, fast sodium 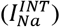 and delayed rectifier potassium currents 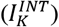. Network connectivity was established through an excitatory glutamatergic synaptic 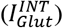 between the pyramidal cell and the interneuron and a GABA-ergic inhibition (*I_GABA_*) from the interneuron to the pyramidal cell. In addition, the pyramidal cell possessed an autoexcitatory glutamatergic synapse (*I_Glut_PC__*) to account for excitatory influences from other pyramidal cells.

In the 2PI-C models, where ion concentrations did not influence the interneuron, intracellular ion concentrations were only calculated for the pyramidal cell. In the extracellular space, the pyramidal cell influenced sodium, potassium, and chloride ions, while the interneuron only contributed to extracellular potassium through its potassium channel. The extracellular space was additionally equipped with a potassium ion sink to account for diffusion and removal of extracellular potassium. The accumulation of ions inside and outside of the pyramidal cell then determined the equilibrium potentials of these ions for the pyramidal cell, while for the interneuron, intracellular ion concentrations and equilibrium potentials were assumed to remain constant (as in previous publications (Desroches et al., 2019)).

In contrast, calculations performed with the 2PI model included the influence of changing ion concentrations on the interneuron and a more complete reciprocal influence of the interneuron on extracellular ion concentrations. In the 2PI models, changes in intracellular ion concentrations were calculated for both neurons, and the interneuron also contained a sodium-potassium pump 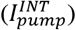. Accordingly, equilibrium potentials were no longer constant for the interneuron in these models and extracellular ion concentrations were thus influenced by both the pyramidal cell and the interneuron.

### 3.1 Interneuron Effects Alter Microcircuit Stability

To test the role of the ion concentrations on the interneuron, and consequently on spike block in the pyramidal cell, we measured the time at which spike block occurred and compared the results for the 2PI and 2PI-C models. Because neuronal excitability and synaptic inhibition have been implicated in spike block initiation, we examined models over a range of neuron excitability and feedback strength. Neuron excitability was varied through depolarizing current injections into the pyramidal cell and interneuron (*I_pc_* and *I_INT_*, respectively), and feedback strength was adjusted by changing GABA maximal conductance (*g_GABA_*). In more stable networks, spike block occurred at a later time, or not at all. Figure 2 shows the comparison of the two models, with cooler colors representing later times of spike block occurrence and thus greater stability. The left column shows the model without ion concentration influence on the interneuron (2PI-C) and the right column shows the model where the interneuron ion concentration effects were included (2PI). As is obvious from the color distribution, including the influence of ion concentrations on the interneuron dramatically stabilized the network. Overall, there were fewer occurrences of spike block in the 2PI model (Figure 2, compare white spaces between models), and when spike block did occur, it was only at high pyramidal cell excitation. In addition, spike blocks occurred later in the 2PI model than in the 2PI-C model (comparatively more cooler colors).

**Fig. 2.**
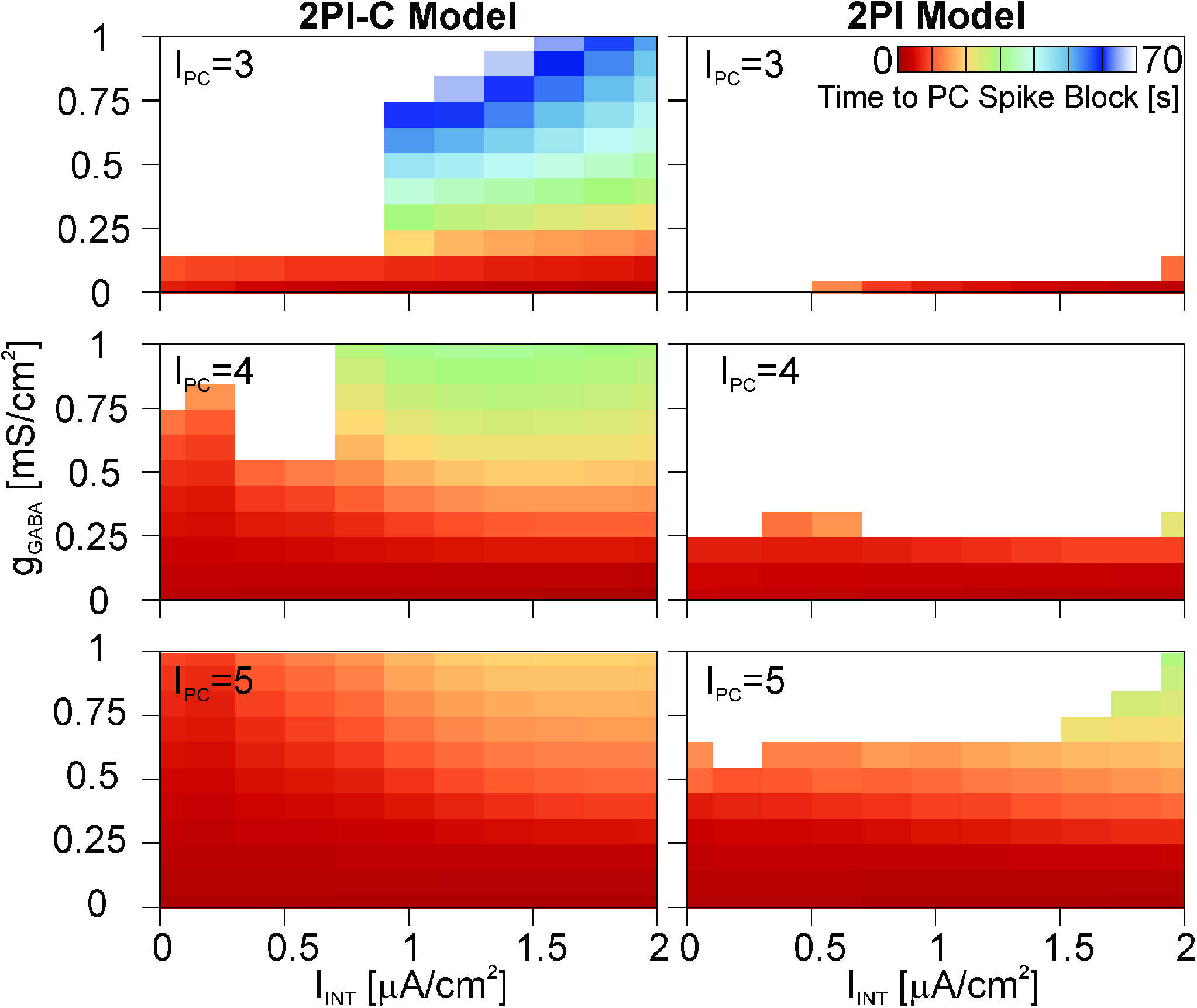
Comparison of models without influence of ion concentrations on the interneuron (2PI-C model, left column) and models where the interneuron influenced, and was affected by, changing ion concentrations (2PI model, right column). White color indicates that no spike block was observed in the pyramidal cell. The colors indicate the time at which spike block occurred in the simulation, with cooler colors representing larger times. Model results are shown as a function of g_GABA_, I_INT_, and I_PC_. Each row is a different *I_pc_* value (μA/cm^2^). *I_pc_* values below 3μA/cm^2^ did not elicit spike block and are not shown. The horizontal axis indicates the excitability of the interneuron via its injected current (*I_INT_*), and the vertical axis shows the synaptic inhibition strength *g_GABA_*.

Figure 2 shows several other notable trends, including more occurrences of spike block with increasing pyramidal cell excitability (*I_pc_*) in both models. This was expected because spike block is initiated by overexcitation and subsequent sodium channel inactivation in the pyramidal cell, making a more excited pyramidal cell inherently less stable against spike block.

Secondly, increasing the GABA inhibition between interneuron and pyramidal cell reduced or delayed spike block occurrence. This was also expected as increased inhibition reduces susceptibility to overexcitation in the pyramidal cell. While these trends were obvious in both models, there were also stark differences between the two models. Importantly, at high pyramidal cell excitability, increasing the injected current into the interneuron (*I_INT_*) had opposing effects on spike block in the pyramidal cell. In the 2PI-C model (left column, Figure 2), injecting more current into the interneuron delayed pyramidal spike block, as evident by cooler colors towards the right side of the *I*_PC_ = 4, 5 panels. However, in the 2PI model (right column, Figure 2), a larger interneuron injected current led to more and earlier instances of spike block. An intuitive explanation for this is that higher firing frequencies of the interneuron enable stronger feedback inhibition, similar to a strengthening of the GABAergic synapse. When the ion concentration effects on the interneuron are neglected, the interneuron’s rapid firing is the dominant mechanism influencing stabilization of the circuit and increased interneuron excitability stabilizes the system.

However, a more active interneuron also facilitates ion concentration changes, and when these effects are included for the interneuron, the behavior of the circuit is altered (right column, Figure 2). In particular, the propensity for spike block increased with high values of interneuron injected current. Therefore, strongly exciting the interneuron helped stabilize the circuit when ion concentrations were excluded from affecting the interneuron but destabilized the circuit when they were allowed to influence the interneuron. This suggests that the interneuron may play a different role in the two models.

### 3.2 Ion Concentration Effects Alter Spike Block Initiation Mechanisms

To determine the mechanism leading to spike block and the role the interneuron might play, we examined the spike activity of the interneuron in more detail. In particular, we found that in the 2PI-C model, when the pyramidal cell exhibited spike block, the interneuron did not (Figure 3A). Instead, in nearly all cases the interneuron continued to produce action potentials. A few rare exceptions were observed when *g_GABA_* was zero, i.e. when the synaptic inhibition between interneuron and pyramidal cell was absent. In these cases, the interneuron stopped firing, but did not exhibit signs of spike block. Unlike during spike block, in these cases, the membrane potential returned to the resting potential (Figure 3B), which indicated that lack of firing was due to the loss of excitation from the pyramidal cell and not an inactivation of the interneuron’s sodium channels. Further evidence that a loss of excitation is responsible for the end of the interneuron’s spiking in these cases can be seen in the inset of Figure 3B, where the interneuron stopped spiking only after the amplitudes of the of the pyramidal cell action potentials fell below 0 mV and thus below transmitter release threshold.

**Fig. 3.**
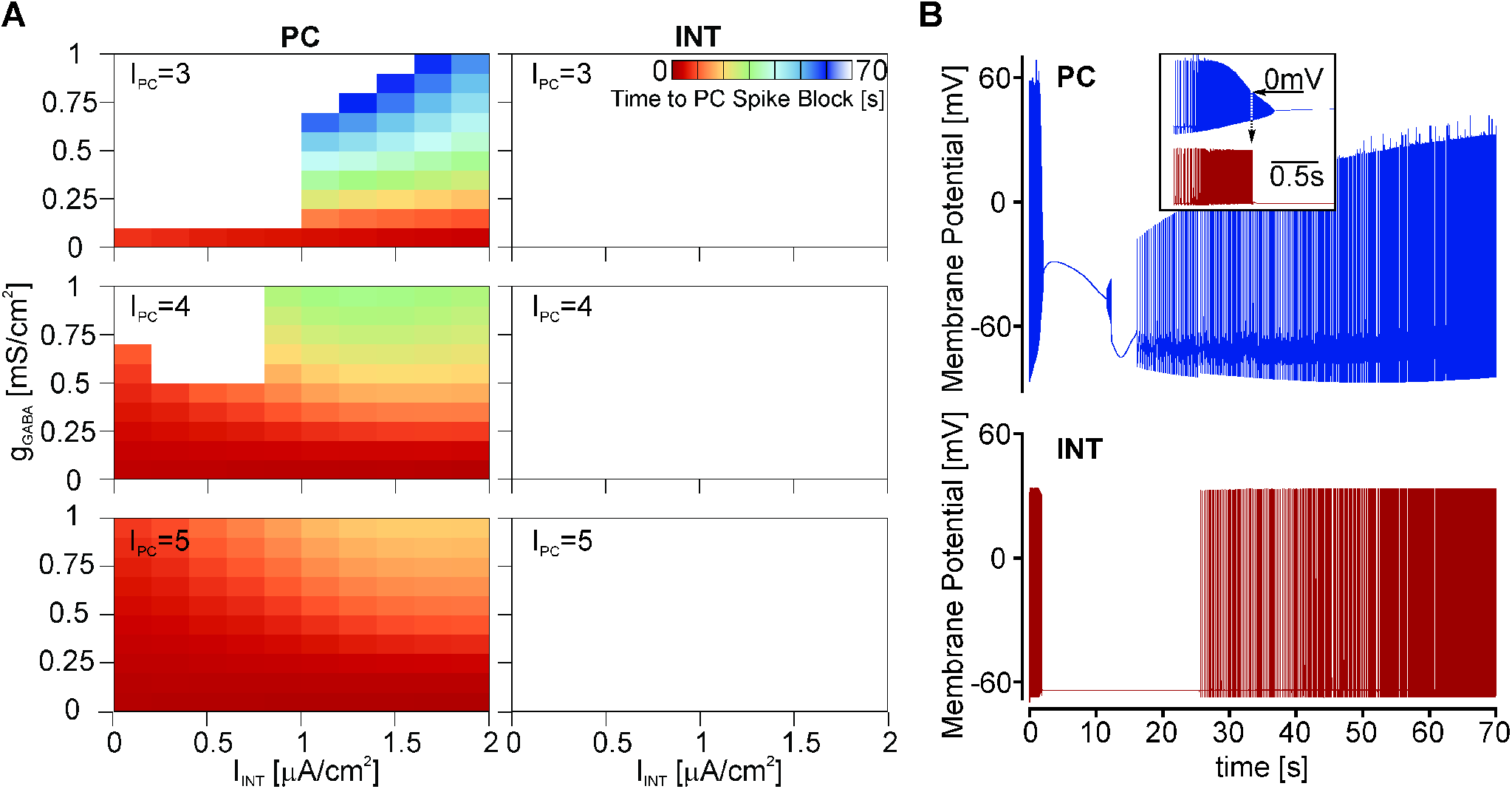
A. In models without influence of ion concentrations on the interneuron (2PI-C model), the interneuron (right column) never showed spike block, despite the frequent occurrence of spike block in the pyramidal cell (left column). The colors indicate the time at which spike block occurred in the simulation, with cooler colors representing larger times. White: no spike block. Model results are shown as a function of *g_GABA_, I_INT_* and *I_pc_* and each row shows a different *I_pc_* value (μA/cm^2^). *I_pc_* values below 3μA/cm^2^ did not elicit spike block and are not shown. The horizontal axis indicates the excitability of the interneuron via its injected current (*I_INT_*), and the vertical axis shows the synaptic inhibition strength *g_GABA_*. B. Example membrane potentials from the pyramidal cell and interneuron during spike block in the pyramidal cell (*I_pc_* = 5 μA/cm^2^, *I_INT_* = 0 μA/cm^2^, *g_GABA_* = 0 mS/cm^2^). The interneuron stopped producing action potentials when the pyramidal spike block occurred. This was due to a loss of glutamatergic excitation from the pyramidal cell and not due to spike block in the interneuron itself. Inset: Magnification, showing that the interneuron stopped firing when the action potential amplitude of the pyramidal cell fell below 0.

In contrast, Fig. 4 shows that in the 2PI model, when ion concentrations were implemented for the interneuron, spike block in the pyramidal cell was always accompanied by spike block in the interneuron. This is obvious from Figure 4A, where the color plots of the pyramidal cell and interneuron are nearly identical. Figure 4B shows the membrane potentials for the same model parameters as in Figure 3B (*I_pc_* = 5 μA/cm^2^, *I_INT_* = 0 μA/cm^2^, *g_GABA_* = 0 mS/cm^2^), and it can be seen that the loss of interneuronal firing was qualitatively very different from that of the 2PI-C model (Figure 3B). Instead of the interneuron membrane potential returning to its resting potential when the pyramidal cell stopped firing, it remained at depolarized values for several seconds before hyperpolarizing below the resting potential, characteristic of sodium channel inactivation and spike block.

**Fig. 4.**
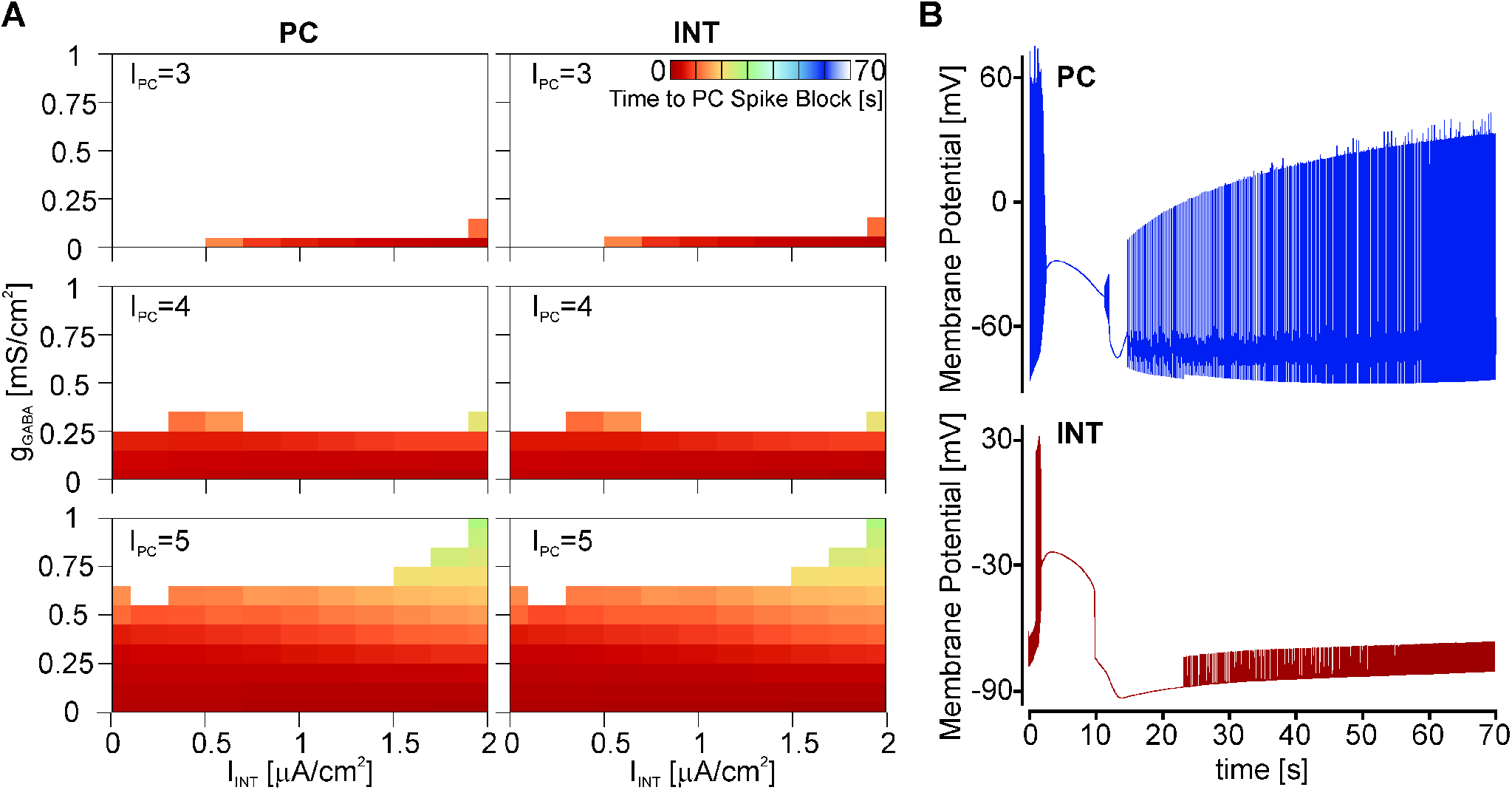
A. In models where ion concentrations affected the interneuron (2PI model), spike block in the interneuron was always observed concurrently with the pyramidal cell. The colors indicate the time at which spike block occurred in the simulation, with cooler colors representing larger times. White: no spike block. Model results are shown as a function of *g_GABA_, I_INT_* and *I_pc_* and each row shows a different *I_pc_* value (μA/cm^2^). *I_pc_* values below 3μA/cm^2^ did not elicit spike block and are not shown. The horizontal axis indicates the excitability of the interneuron via its injected current (*I_INT_*), and the vertical axis shows the synaptic inhibition strength *g_GABA_*. B. Example membrane potentials from the pyramidal cell and interneuron during spike block in the pyramidal cell (*I_pc_* = 5 μA/cm^2^, *I_INT_* = 0 μA/cm^2^, *g_GABA_* = 0 mS/cm^2^). Both the pyramidal cell and the interneuron exhibit spike block, with spike block in the pyramidal cell occurring shortly after spike block in the interneuron.

Most strikingly, the interneuron spike block always preceded the pyramidal cell spike block. This can be seen in the selected example recordings in Figure 5A. Examples were taken from a variety of parameter combinations in the 2PI model that resulted in spike block at various times during the simulation. In Figure 5B, the time difference between spike block occurrence in the pyramidal cell and the interneuron is shown for all parameter combinations. Specifically, we found that when spike block occurred, the interneuron always preceded the pyramidal cell by 0.2 - 0.4 seconds. We observed that with more current injection into the interneuron (*I_INT_*), the delay between interneuron spike block and the subsequent pyramidal cell spike block increased (warmer colors to the right in Figure 5B). This is consistent with higher interneuron firing frequencies providing increasing inhibition to the pyramidal cell, leading to lower pyramidal cell firing frequencies and longer times to spike block. We also noted that with higher pyramidal cell excitation (*I_pc_*), the delay between interneuron spike block and the subsequent pyramidal cell spike block was shorter (fewer warmer colors in the bottom panel of Figure 5B). This was a result of the pyramidal cell being more excited and therefore closer to spike block.

**Fig. 5.**
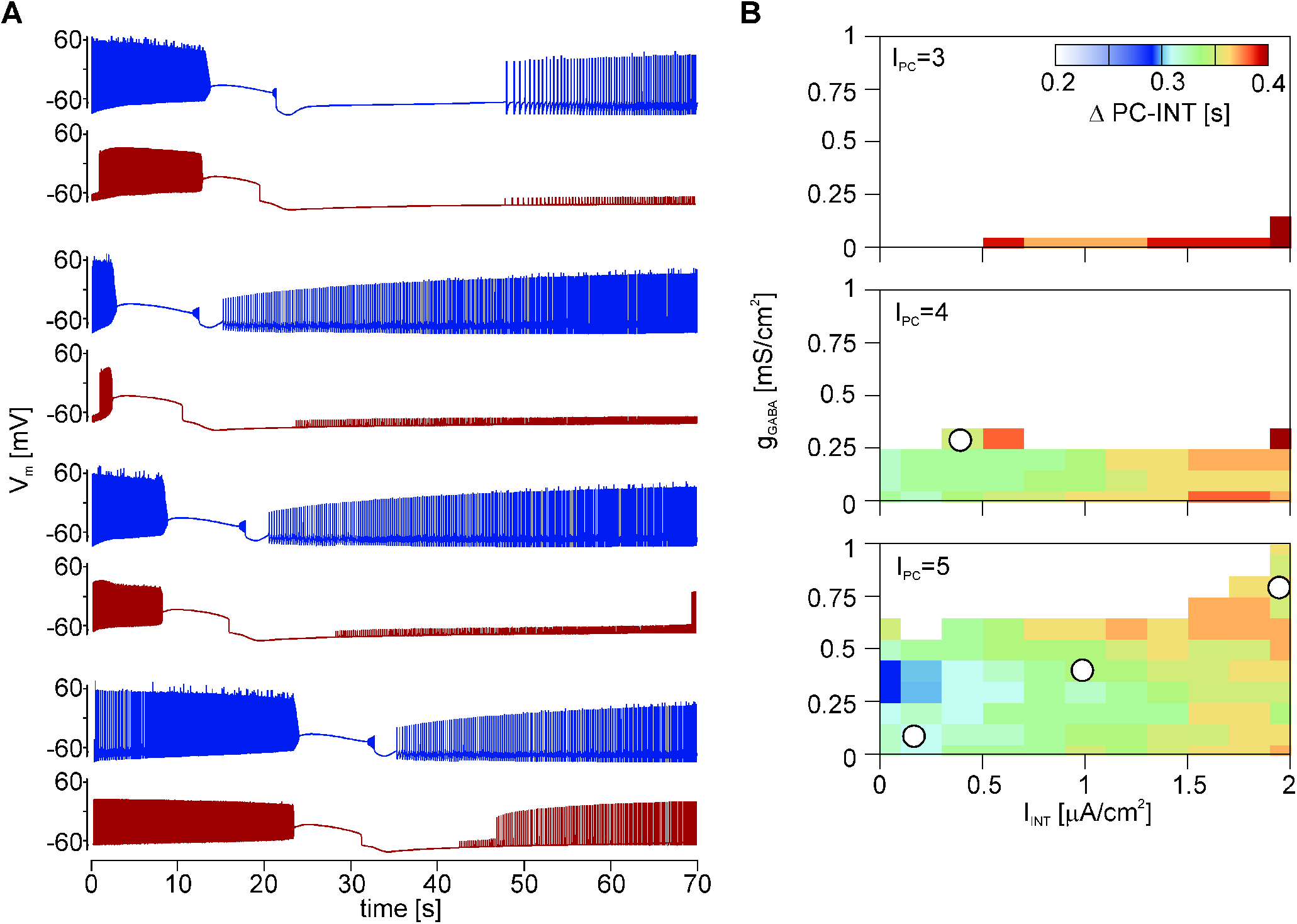
A. Membrane potentials of the 2PI model for the interneuron (red) and the pyramidal cell (blue) as a function of time for four cases that show spike block at different times during the simulation (from top to bottom: *I_PC_* = 4 μA/cm^2^, *I_INT_* = 0.4 μA/cm^2^, *g_GABA_* = 0.3 mS/cm^2^; *I_pc_* = 5 μA/cm^2^, *I_INT_* = 0.2 μA/cm^2^, *g_GABA_* = 0.1 mS/cm^2^;. *I_pc_* = 5 μA/cm^2^, *I_INT_* = 1.0 μA/cm^2^, *g_GABA_* = 0.4 mS/cm^2^; *I_pc_* = 5 μA/cm^2^, *I_INT_* = 2.0 μA/cm^2^, *g_GABA_* = 0.8 mS/cm^2^); model parameters are highlighted with dots in figure 5B, as well. In all cases, spike block in the interneuron preceded spike block in the pyramidal cell. B. Time delay (Δ) between the occurrence of spike block in the interneuron and the pyramidal cell. Positive numbers indicate that spike block occurred first in the interneuron and cooler colors indicate shorter delays. Highlighted dots: Example models shown in A. *I_pc_* values are given in μA/cm^2^.

Overall, our data indicate that the mechanism leading to spike block differed depending on whether or not the interneuron influenced, and was influenced by, the ion concentrations. Without this influence, spike block initiation was correlated primarily with pyramidal cell excitability. In contrast, with the influence of ion concentrations, the interneuron appeared to play a critical role in eliciting the pyramidal cell’s spike block.

### 3.3 Spike Block in the Interneuron Triggers Spike Block in the Pyramidal Cell

To test the critical impact of the interneuron and the hypothesis that indeed the loss of interneuronal spiking elicited the spike block in the pyramidal cell, we manipulated the interneuron in the 2PI model to either stop firing prematurely or to continue firing indefinitely. If the interneuronal spike block elicits the pyramidal cell spike block, then prematurely stopping interneuronal firing should cause the pyramidal cell to also exhibit spike block prematurely. Complementary to this, extending interneuronal firing should prevent or delay the pyramidal cell’s spike block.

Figure 6 shows results from simulations (*I_pc_* = 5 μA/cm^2^, *I_INT_* = 1 μA/cm^2^, *g_GABA_* = 0.4 mS/cm^2^) where we prematurely ended interneuronal firing by setting the interneuron membrane potential to a constant value. In the left column, the membrane potential was set to −25mV, mimicking the membrane potential when spike block occurs (compare to Fig. 5A). In the right column, it was set to −70mV, i.e. the resting membrane potential, corresponding to a loss of interneuronal spiking without spike block (compare to Figure 3B). Our results show that when the interneuron ceased firing, it triggered spike block in the pyramidal cell, independent of the interneuronal membrane potential value.

**Fig. 6.**
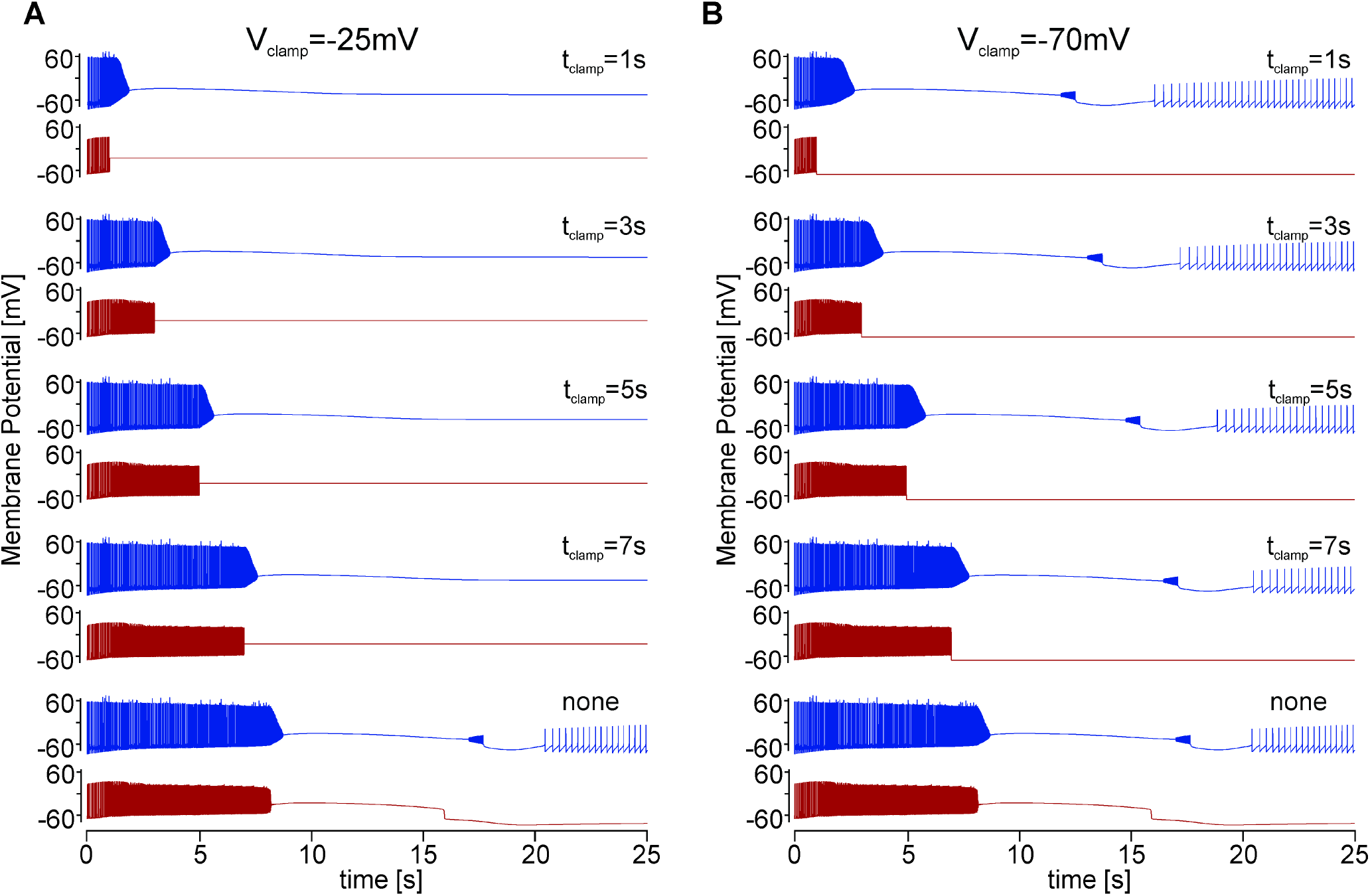
Prematurely ending interneuronal spiking elicits spike block in the pyramidal cell. Membrane potentials of the interneuron (red) and the pyramidal cell (blue) as a function of time. The bottom recordings (labeled “none”) in each column show the spontaneously occurring spike blocks with the following parameters: *I_pc_* = 5 μA/cm^2^, *I_INT_* = 1 μA/cm^2^, *g_GABA_* = 0.4 mS/cm^2^. The other recordings show simulation results where the membrane potential of the interneuron was clamped to a fixed value (labeled with *t_clamp_*), at increasingly later times during the simulation (top to bottom). In the left column, the interneuron membrane potential was clamped to −25 mV, in the right column to −70 mV.

We observed that when the interneuron was stopped prematurely, the spike block in the pyramidal cell also occurred prematurely, and that earlier interneuron stop times resulted in earlier spike block times of the pyramidal cell. In all cases, the pyramidal cell spike block occurred shortly after firing ceased in the interneuron. We noted that the time delay between the two events increased with earlier interneuronal stop times, as seen in Figures 6 and 7A. For example, when the interneuron membrane potential was set to −25mV at 6s, the delay was 0.40s, while the same intervention at 1s resulted in a delay of 0.68s.

**Fig. 7.**
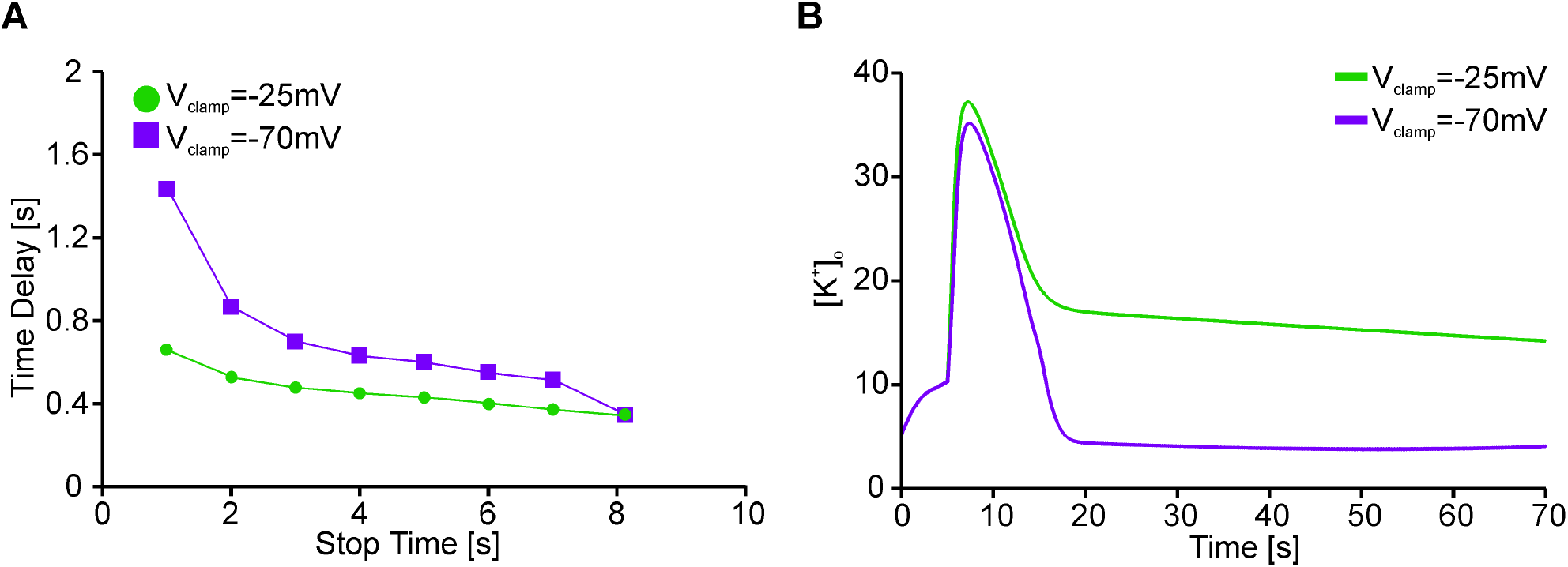
A. Time delay between spike block occurrence in the pyramidal cell and the end of spiking in the interneuron as a function of when interneuronal spiking was stopped. The examples shown come from models with the following parameters: *I_pc_* = 5 μA/cm^2^, *I_INT_* = 1 μA/cm^2^, *g_GABA_* = 0.4 mS/cm^2^. Without manipulation, these parameters induce spike block in both the interneuron and the pyramidal cell (compare figure 6, bottom). Earlier stop times resulted in longer delays, independently of whether spiking was stopped by clamping the interneuron membrane potential to −25 mV (green) or −70 mV (purple). Note that both curves converge on the same data point at 8.15 seconds because this is the unperturbed condition when spike block occurred without forced changes to the interneuron membrane potential. B. Changes of the extracellular potassium concentrations when interneuronal firing was stopped at 5 s (green: −25 mV clamping potential, purple: −70 mV).

The time delay results in Fig. 7A were qualitatively independent of whether the interneuron was held at −25mV or −70mV, although the quantitative values differed. The longer time delays for an interneuron membrane potential clamped at −70 mV compared to −25 mV were likely due to less activation of voltage-gated ion channels in the interneuron and thus slower ion concentration changes in the extracellular space. Indeed, when we looked at the accumulation of extracellular potassium in the two conditions (Figure 7B), there was a smaller and less sustained increase of extracellular potassium when the interneuron was held at −70 mV (purple line). The increased delays at −70 mV were thus likely caused by less accumulation of extracellular potassium, which decreased the excitability of the pyramidal cell on its path towards spike block.

In addition to the time delay differences, we also noted qualitative differences in the behavior of the pyramidal cell after spike block that depended on the interneuron membrane potential value. Specifically, at an interneuronal membrane potential of −70mV, the pyramidal cell started to fire again, while at −25mV it did not. This was also likely due to the differences in extracellular potassium concentrations. At −25mV, there was a sustained potassium leak from the interneuron, causing a continuous excitation of the pyramidal cell. Consequently, the pyramidal cell sodium channels were unable to de-inactivate and remained inactivated after the spike block, which was characterized by a depolarized membrane potential without action potentials (Fig. 6A). In contrast, when the interneuron membrane potential was held at −70 mV, less potassium was released into the extracellular space (Figure 7B, green line) and the pyramidal cell was able to recover from spike block. Recovery was possible because the pyramidal cell membrane potential hyperpolarized rapidly about 10 seconds after spike block (Figure 6B), allowing sodium channels to de-inactivate and new action potentials to eventually begin again. Together, the above simulations suggest that spike block in the interneuron is responsible for triggering spike block in the pyramidal cell. Furthermore, this chain of events is likely mediated through rapid changes in extracellular ion concentrations.

Additional evidence that the loss of interneuronal inhibition and sharp changes in ion concentrations play critical roles initiating pyramidal cell spike block comes from simulations in which the firing of the interneuron was artificially extended. Figure 8A (left column) shows the membrane potentials for the same parameter combinations as used in Figure 6 (*I_pc_* = 5 μA/cm^2^, *I_INT_* = 1 μA/cm^2^, *g_GABA_* = 0.4 mS/cm^2^). Here, however, instead of stopping interneuronal firing prematurely, it was continued through the end of the simulation. Specifically, the interneuronal membrane potential was stored prior to the occurrence of spike block (from 6 to 6.084 seconds, 10 spikes, 119 Hz) and then repeated indefinitely. Despite the continuous interneuronal activity, the pyramidal cell exhibited spike block, and even at an earlier time (7.84s) than under normal conditions (8.15s). Thus, while a loss of inhibition from the interneuron prematurely triggered spike block (Figures 6 and 7), preventing this loss did not stabilize the pyramidal cell against spike block. Indeed, when we looked at the changes in extracellular ion concentrations, we found that they mirrored those in the unperturbed conditions. Specifically, the extracellular sodium concentration dropped dramatically during spike block, while the extracellular potassium concentration increased. This happened regardless of whether or not the interneuronal firing was maintained.

**Fig. 8.**
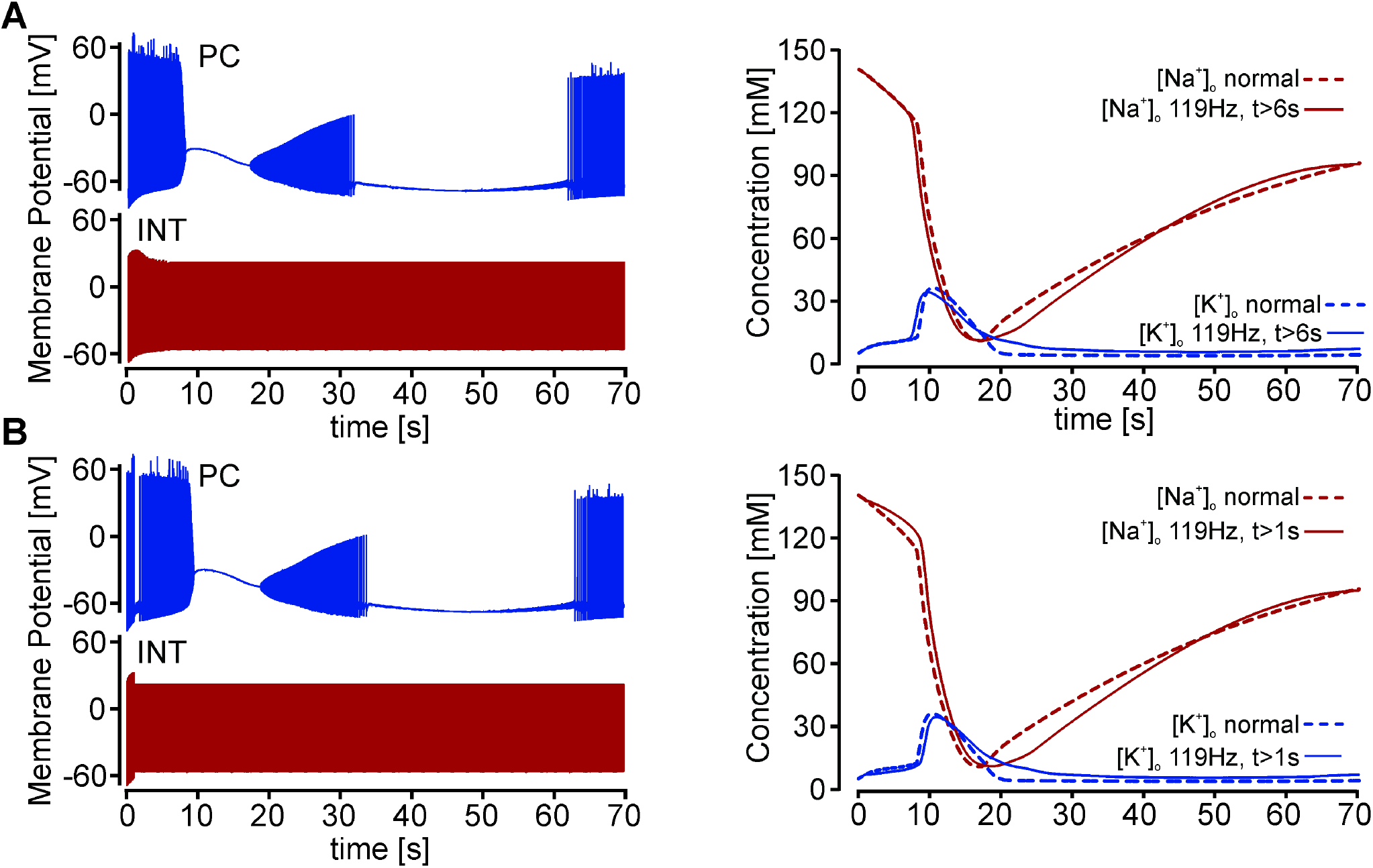
Left: Membrane potentials of the interneuron (red) and the pyramidal cell (blue) as a function of time. Right: Extracellular sodium (red) and potassium (blue) over time. The examples are taken from a model with the following parameters: *I_pc_* = 5 μA/cm^2^, *I_INT_* = 1 μA/cm^2^, *g_GABA_* = 0.4 mS/cm^2^. Without manipulation, these parameters induce spike block in the both neurons (compare figure 6, bottom). A. Interneuron firing frequency was set to 119 Hz at 6 seconds and repeated indefinitely until the end of the simulation. This did not prevent spike block in the pyramidal cell. Changes in extracellular ion concentrations (solid lines) mirrored those in unperturbed conditions (‘normal’, dashed lines). B. Interneuron firing frequency was set to 119 Hz at 1 second and repeated indefinitely until the end of the simulation. This did not prevent spike block in the pyramidal cell. Changes in extracellular ion concentrations mirrored those in unperturbed conditions.

The results from Figures 6-8 indicated that the extracellular ion concentrations mediated the occurrence of spike block in the pyramidal cell. Therefore, it is possible that at 6s into the simulation, the pyramidal cell was already on an unavoidable path to spike block and the sustained inhibition from the interneuron was insufficient to prevent spike block.

To test whether the buildup of ion concentrations could be reduced, and thus spike block prevented, we repeated the simulation of Figure 8A but held the interneuron firing frequency constant (119 Hz) from an earlier point in time (1 s). The hypothesis was that this earlier intervention could interrupt the concentration changes that lead to spike block. We found that earlier intervention delayed pyramidal cell spike block (9.18 s), but did not prevent it. This suggested that the occurrence of the pyramidal cell spike block does not always require interneuron spike block, but rather can be initiated through rapid changes in ion concentrations.

### 3.4 High Firing Frequencies lead to Elevated Extracellular Potassium that Elicit Spike Block

As a proof of concept that changes in extracellular ion concentrations are sufficient to elicit pyramidal cell spike block, we took the time-dependent ion concentrations from a model that normally exhibits spike block (*I_pc_* = 5 μA/cm^2^, *I_INT_* = 1 μA/cm^2^, *g_GABA_* = 0.4 mS/cm^2^) and implemented them in a model that does not normally exhibit spike block (*I_pc_* = 3 μA/cm^2^, *I_INT_* = 1 μA/cm^2^, *g_GABA_* = 0.4 mS/cm^2^). All other model parameters were allowed to evolve freely. Figure 9 shows the ion concentrations over time for both sets of model parameters. Stark differences in extracellular sodium and potassium concentrations are apparent. In the model that does not normally show spike block, the extracellular sodium and potassium concentrations remained relatively constant (dashed lines). However, when spike block did occur, it was accompanied by rapid changes in both sodium and potassium concentrations (solid lines).

**Fig. 9.**
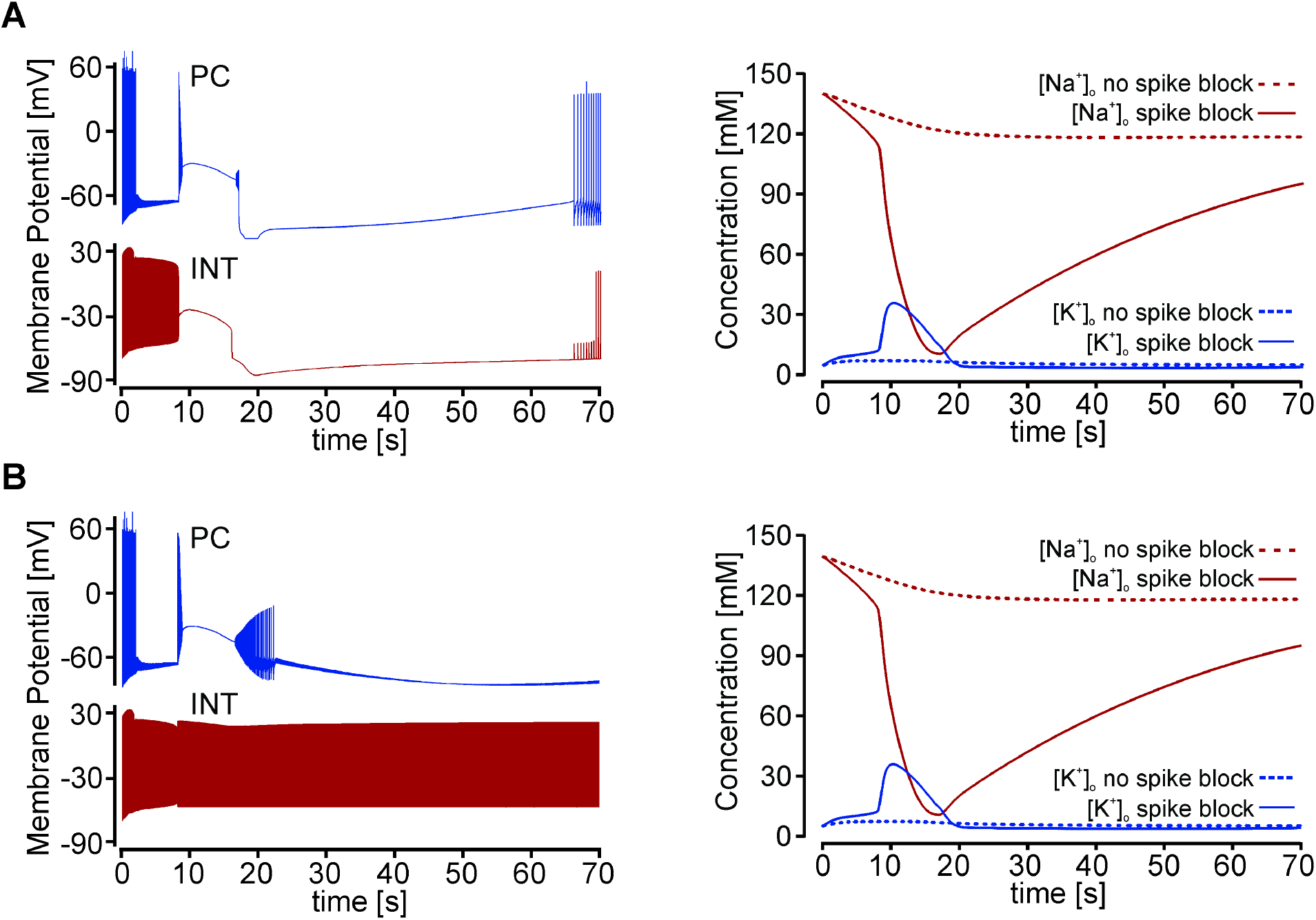
Left: Membrane potentials of the interneuron (red) and the pyramidal cell (blue) as a function of time. Right: Extracellular sodium (red) and potassium (blue) concentrations over time. The dashed curves are the concentrations from a model that does not normally exhibit spike block in either neuron with the parameters *I_pc_* = 3 μA/cm^2^, *I_INT_* = 1 μA/cm^2^, *g_GABA_* = 0.4 mS/cm^2^. The solid curves are the concentrations from a model that normally exhibits spike block with the parameters *I_pc_* = 5 μA/cm^2^, *I_INT_* = 1 μA/cm^2^, *g_GABA_* = 0.4 mS/cm^2^ (compare to figure 6 bottom). A. With no other manipulation, forcing the extracellular ion concentrations in the model that does not normally exhibit spike block to match those of the model that does show spike block was sufficient to elicit spike block in both the interneuron and the pyramidal cell. B. Same model as in A, but here the interneuron was additionally prevented from spike block by fixing its firing frequency to 119 Hz starting at 8.15 seconds and maintaining it throughout the simulation. Preventing spike block in the interneuron did not eliminate spike block in the pyramidal cell.

Figure 9A shows that the implementation of rapid ion concentration changes can elicit interneuron and pyramidal cell spike block in a model that does not normally show it. Like in previous models, the interneuron’s spike block preceded that of the pyramidal cell. To separate the effects of loss of firing in the interneuron (and a subsequent loss of inhibition of the pyramidal cell) from ion concentration changes on the pyramidal cell, we then repeated the simulation, but with the interneuronal firing frequency maintained at 119 Hz beginning at the unperturbed spike block time. Figure 9B shows that pyramidal cell spike block occurred even when the interneuron was prevented from spike block. Taken together, the simulations in Fig. 9 demonstrated that changes in extracellular ion concentrations are sufficient to induce spike block in the pyramidal cell.

To determine whether rapid changes in extracellular ion concentrations are not simply sufficient, but necessary to elicit spike block, we fixed the extracellular ion concentrations at their values shortly before spike block (t = 6 s) in the model that naturally shows spike block (same as Figures 6 and 8, *I_pc_* = 5 μA/cm^2^, *I_INT_* = 1 μA/cm^2^, *g_GABA_* = 0.4 mS/cm^2^). Figure 10A shows that holding the extracellular concentrations of both sodium and potassium at their pre-spike block values prevented spike block. The interneuron still exhibited high firing frequencies near 100 Hz, however the pyramidal cell firing frequency remained low near 40 Hz.

**Fig. 10.**
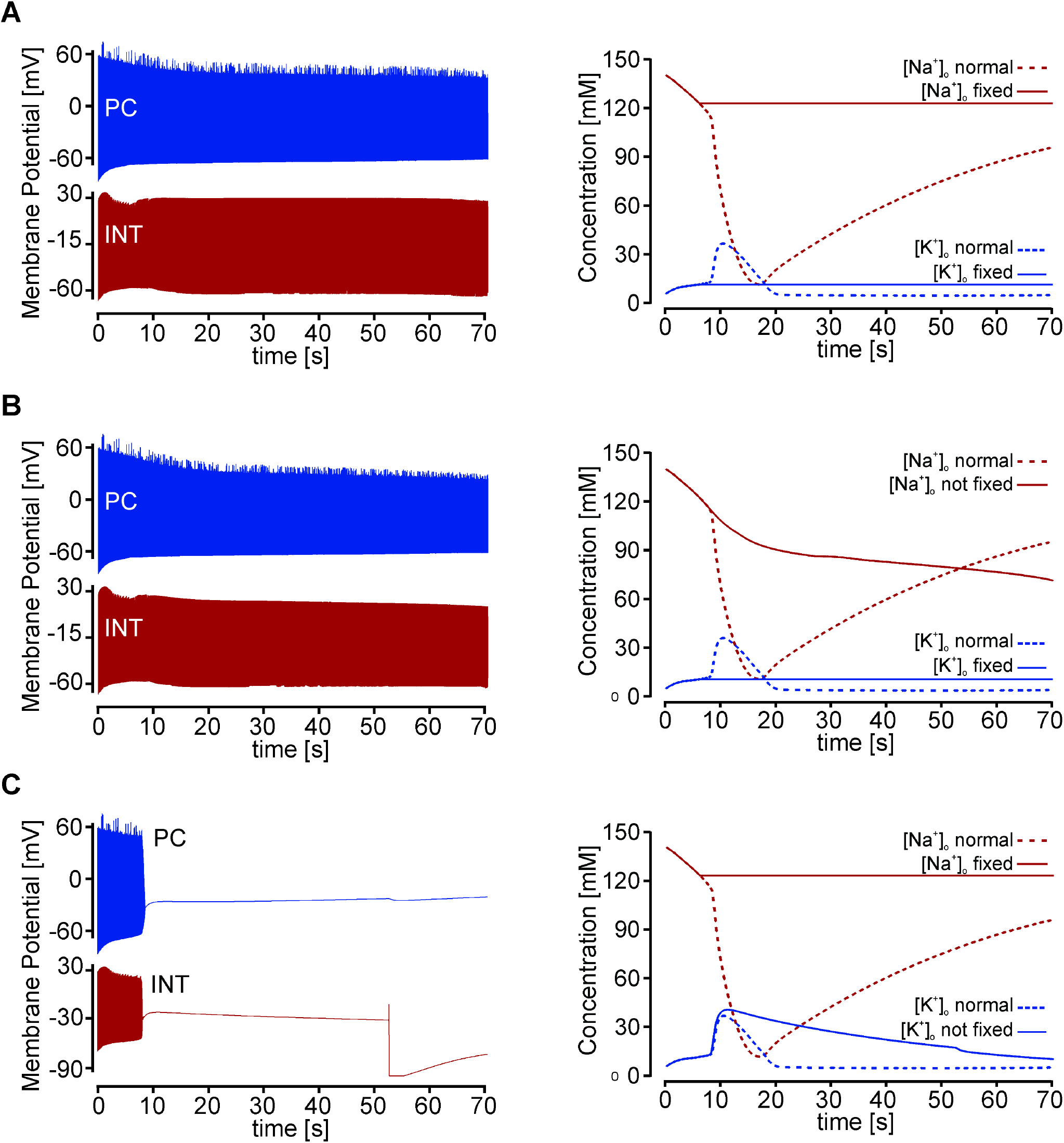
Extracellular potassium concentration changes are necessary to elicit spike block. Left: Membrane potentials of the interneuron (red) and the pyramidal cell (blue) as a function of time. Right: Extracellular sodium (red) and potassium (blue) concentrations over time. The examples are taken from a model with the following parameters: *I_pc_* = 5 μA/cm^2^, *I_INT_* = 1 μA/cm^2^, *g_CABA_* = 0.4 mS/cm^2^. Without manipulation, these parameters induce spike block in both neurons. A. When extracellular ion concentrations were fixed to 10.7 mM (potassium) and 122.7 mM (sodium) after 6 seconds, spike block did not occur B. Spike block was still prevented when only the potassium concentration was held at a fixed value (10.7 mM). C. When only the sodium concentration was held constant (122.7 mM), spike block occurred in both neurons.

To identify whether both sodium and potassium concentration changes, or only one, were necessary to induce spike block in the pyramidal cell, we also ran simulations in which only a single extracellular ion concentration was fixed. Figure 10B shows that when solely the potassium concentration was held constant, spike block in the pyramidal cell was prevented. Again, the interneuron exhibited high firing frequencies while the pyramidal cell firing frequency remained low. In contrast, when only the sodium concentration was held constant (Figure 10C), pyramidal cell spike block was observed. In this case, the interneuronal and pyramidal firing frequencies increased dramatically before both neurons stopped firing (200 to 600 Hz), as seen in other models with spike block. This suggested that high interneuronal firing frequency alone does not cause spike block. Rather, the increase in extracellular potassium concentration caused by the high firing frequency is critical to elicit spike block.

In the cases above, the prevention of potassium concentration changes was accompanied by continued spiking of the interneuron. It was thus not clear whether the continued interneuronal spiking and its continued inhibition of the pyramidal cell contributed to preventing the spike block in the pyramidal cell. Therefore, we repeated the simulation from Figure 10B, keeping the potassium concentration constant while additionally stopping interneuronal firing at t = 6 s. The pyramidal cell did not exhibit spike block, but a brief, rapid increase in firing frequency to more than 200 Hz was observed. The rapid firing lasted for a few seconds before the pyramidal cell settled into a steady firing rate of approximately 70 Hz. This is further indication that while a loss of inhibition leads to increased firing frequencies of the pyramidal cell, it is not sufficient to elicit spike block. Instead, ion concentration changes are essential to eliciting spike block in the pyramidal cell.

Overall, we find that in all models, the interneuron is key to the initiation of the pyramidal cell spike block. The inhibition the interneuron exerts on the pyramidal cell partially protects the pyramidal cell from spike block. However, the high firing frequencies of both neurons, combined with rapid changes in extracellular ion concentrations, initiate spike block in the interneuron. This sudden loss of inhibition results in the pyramidal cell also exhibiting spike block. Thus, interneuronal firing stabilizes cortical microcircuits, but its loss can destabilize the system and result in depression of the entire circuit.

## 4 Discussion

Despite widespread prevalence, the root cause of migraine, CSD, and the neuronal predispositions that facilitate them remain largely unknown. In a small subset of people with familial hemiplegic migraine (FHM), migraine attacks occur with aura and hemiparesis (Tiwari et al., 2020). While relatively rare, identification of the genetic origins of FHM has allowed for some of the underlying cellular mechanisms to be identified, which has helped provide clues to the cause of other types of CSD and migraine. Among the underlying causes of FHM are mutations in ion channels and pumps that can destabilize cortical circuits, leading to overexcitation and ultimately a traveling wave of depolarization-induced spike block. The consequences of mutations in ion pumps, such as the loss-of-function mutation in the alpha2 subunit of the glial Na+/K+ pump in FHM-2 (De Fusco et al., 2003) are conceptually easy to understand because of their direct impacts on the dynamics of ion concentration changes. In contrast, the consequences of mutations in ion channels are less well understood, as their pathological mechanisms may depend on the sign (gain of function vs. loss of function), the efficacy of the mutation, and which neurons express the channel. Even in simplified microcircuits, like the one used in this study, the negative feedback between pyramidal cells and interneurons makes predictions for the influence of ion channel mutations on circuit activity difficult.

FHM is not the only family of diseases with known ion channel mutations that affect cortical excitation. Monogenic diseases related to mutations in voltage-gated sodium channels are well documented in patients with epileptic phenotypes (Menezes et al., 2020; Poulin & Chahine, 2021), including almost 2,000 identified mutations that affect sodium channel isoforms highly expressed in brain tissues (including NaV1.1, NaV1.2, NaV1.3 and NaV1.6; (Huang et al., 2017)). One of these mutations (NaV1.1) has also been identified in FHM-3 patients (Tiwari et al., 2020), although its role in migraines and CSD is less clear. In particular, it remains an open question whether the FHM-3 mutation leads to a gain or loss of function of the channel (Hedrich et al., 2014; Mantegazza & Broccoli, 2019). NaV1.1 channels are particularly important for interneuronal excitability, and it is also unclear how changes in this excitability may contribute to initiating CSD. For example, increased activity of GABAergic interneurons through a gain of function mutation should reduce excitability of cortical networks through increased inhibition of the pyramidal cells, while loss-of-function mutations should do the opposite.

In our study, we tested the effects of both increased and decreased excitability of the GABAergic interneurons on spike block occurrence, taking into account changes in ion concentrations that may result from high firing frequencies. Our comparison of models with and without the inclusion of ion concentration effects on the interneuron demonstrated that their inclusion led to a wider dynamic range of the circuit and generally improved stability. Additionally, contrary to expectation, our models showed that reduced interneuronal excitability was always associated with a more stable circuit and decreased tendency for spike block. Consistent with this, we also found that increased interneuron excitability, as predicted for a NaV1.1 gain-of-function mutation, always destabilized circuit behavior. This destabilization occurred despite higher interneuronal firing rates, causing more inhibition of the pyramidal cell. In this case, the interneuron became hyperexcited, furthering the rapid changes in intra- and extracellular ion concentrations that ultimately led to spike block in both the interneuron and the pyramidal cell. Combined, these results are consistent with the hypothesis that the FMH3 NaV1,1 mutation is a gain of function mutation.

The complexity of ion channel mutations on circuit behavior is also apparent when the feedback between the pyramidal cell and the interneuron is considered. In general, hyperexcitability of pyramidal cells predisposes them to spike block, but this overexcitation also increases the excitation of the interneurons, which in turn provides greater inhibition to the pyramidal cell. Indeed, our model suggests that this inhibition is crucial to the stability of the circuit because when it was lost, pyramidal cell spike block was always initiated. However, if the interneuronal inhibition was maintained, this did not rescue the pyramidal cell from spike block. In fact, we found that increasing interneuronal excitation even facilitated instability in the circuit and made spike block more likely to occur.

The effect of the interneuron on the stability of the circuit is thus multifaceted, with high or maintained firing frequencies accumulating potassium in the extracellular space, which facilitates spike block. At the same time, high interneuronal firing frequencies increase GABA release, which our data clearly demonstrate stabilizes the circuit by diminishing pyramidal cell excitation. Pathologies of the GABA synaptic strength or their temporal dynamics, like facilitation or depression, may thus play into initiating, or preventing, spike block in pyramidal cells. Interneuronal inhibition of the pyramidal cells is exerted through ionotropic GABA-A receptors, although evidence is mounting that paracrine GABA release may additionally act on GABA-B receptors (Kulik et al., 2018). Interestingly though, pharmacological approaches using GABA agonists such as the anticonvulsant diazepam to supernaturally activate GABA receptors have failed to provide clear evidence for reducing CSD and migraines (reviewed in (C. Ayata, 2009)). Clearly there are many competing factors when broadly applying drugs, but our model also gave an indication for why such GABA agonists may fail to alleviate CSD. In our model, an increase in GABA actions was implemented by increasing the maximum synaptic conductance. This, however, is equivalent to adding additional GABA receptors, which increases GABA actions, and with it the inhibition of the pyramidal cell, by several fold. In contrast, GABA-A agonists will only maximally activate existing GABA receptors, which may not lead to a sufficient increase in GABAergic inhibition. Indeed, when we maximally activated the GABAergic synapse between interneuron and pyramidal cell, we found that there was only a mild effect on the occurrence of spike block (Supplemental figure 1). At high excitability levels of the pyramidal cell, maximally activating the GABAergic synapse was unable to prevent spike block. These observations are consistent with experimental results that show that potent anticonvulsants failed to prevent CSD, and may even explain why in some cases extreme pentobarbital doses increased the threshold for CSD (Van Harreveld & Stamm, 1953).

Regardless of the underlying mechanism, in all our models, the universal driving condition leading to spike block was a rapid change in extracellular ion concentrations. Our results showed that these ion concentration changes alone were sufficient to initiate spike block and that they are a result of high frequency firing of both cells, which initiates spike block in the interneuron and subsequently the pyramidal cell. The criticality of rapid extracellular ion concentration changes in initiating spike block, in particular a rapid increase of extracellular potassium, has implications beyond the current model. For example, large potassium efflux is known to cause osmotic cell swelling and a concurrent reduction in extracellular space of more than 50% (Cenk Ayata & Lauritzen, 2015). Our models used a conservative estimate for the extracellular volume, and it seems logical to assume that the inclusion of cell swelling will likely serve to accelerate extracellular ion concentration changes and thus also the occurrence of CSD.

While our model focused on a two-cell microcircuit, our findings have implications for a possible mechanism for the initiation of CSD within cortical networks. We have shown that the interneuron is critical to the initiation of pyramidal cell spike block. Within the context of a larger network, where a single interneuron can innervate many pyramidal cells, the loss of spiking in one or a few interneurons could act as a source for the larger scale spreading wave of cortical depression. Thus, our results point to the possibility that localized effects on cortical interneurons can have global implications beyond the local microcircuit.

## Supplementary Information

**Supplemental figure 1.**
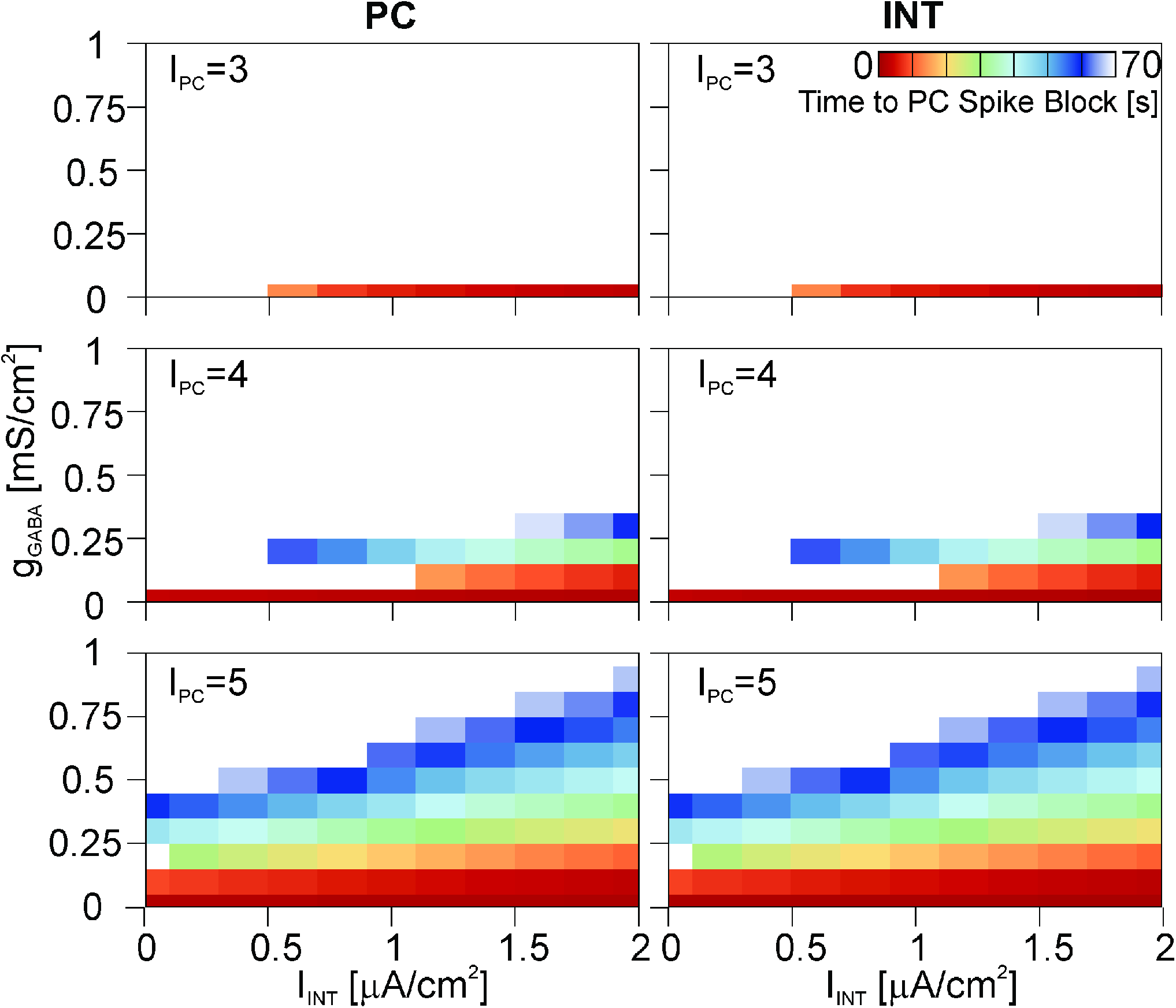
Maximal activation of the GABAergic synapse between interneuron and pyramidal cell (*s_GABA_* = 1) did not prevent spike block from occurring. The colors indicate the time at which spike block occurred in the simulation (left column: pyramidal cell, right column: interneuron), with cooler colors representing larger times. White: no spike block. Model results are shown as a function of *g_GABA_, I_INT_*, and *I_pc_* and each row shows a different *I_pc_* value (μA/cm^2^). *I_pc_* values below 3μA/cm^2^ did not elicit spike block and are not shown. The horizontal axis indicates the excitability of the interneuron via its injected current (*I_INT_*), and the vertical axis shows the synaptic inhibition strength *g_GABA_*.

## Footnotes

1 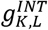 and 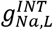 were selected to match a total conductance of 0.1 mS/cm^2^ and a reversal potential of −65 mV (Wang & Buzsáki, 1996)

2 Initial intracellular concentrations were chosen to match the equilibrium potentials.

## Notes

### Competing Interest Statement

The authors have declared no competing interest.

### Summary of Updates

Added link to modelDB database with the source code for the used model. Formatting was made more printer friendly (line space reduced).

